# GlycoGenius: the ultimate high-throughput glycan composition identification tool

**DOI:** 10.1101/2025.03.10.642485

**Authors:** Hector F. Loponte, Jing Zheng, Yajie Ding, Isadora A. Oliveira, Kristoffer Basse, Adriane R. Todeschini, Peter L. Horvatovich, Guinevere S.M. Lageveen-Kammeijer

## Abstract

Mass spectrometry is recognized as the gold standard for glycan analysis, yet the complexity of generated data hampers progress in glycobiology, as existing tools lack full automation, requiring extensive manual effort. We introduce GlycoGenius, an open-source program offering an automated workflow for glycomics data analysis, featuring an intuitive graphical interface. With algorithms tailored to reduce manual workload, it allows for data visualization and automatically constructs search spaces, identifies, scores, and quantifies glycans, filters results, and annotates fragment spectra of *N-* and *O-*glycans, glycosaminoglycans and more. It seamlessly guides researchers of all expertise levels from raw data to publication-ready figures. Our findings demonstrate that GlycoGenius achieves results comparable with manual analysis or competing tools, identifying more glycans, including novel ones, while significantly reducing processing time. This groundbreaking tool represents a significant advancement in the study of glycoconjugates, empowering researchers to focus on insights rather than data processing.

## INTRODUCTION

Glycoconjugates play a crucial role in defining cellular interactions and functions, forming an integral part of the cellular environment as secreted molecules or components of the glycocalyx. These complex carbohydrates, encompassing glycoproteins, proteoglycans, and glycolipids, mediate a diverse array of interactions between cells and their surroundings ^1^. The glycosylation patterns of cells impact various cellular functions, including signal transduction, cell-cell and cell-pathogens interactions, adhesion, motility, protein folding and stability, growth regulation, cell differentiation, as well as immune modulation through the activation and inhibition of specific pathways ^2^. This wide range of functions underscores the role of glycoconjugates in maintaining cellular homeostasis and their dysregulation in pathological conditions like diabetes ^3^, cancer ^4^, Alzheimer’s^5^ and Parkinson’s^6^ diseases highlighting their significance in clinical research. Alterations in the glycomic profile may act as potential therapeutic targets^7^ or as diagnostic molecular markers ^8^, emphasizing the urgent need for advanced analytical techniques to decode the structural and functional complexity of glycoconjugates, which would foster deeper insights into their involvement in diverse pathologies and normal physiological processes.

The main analytical platform for in-depth characterization and structure elucidation of glycans is mass spectrometry (MS) ^9^. While full scan MS1 spectra enable the analysis of various compounds within a sample ^10^, fragmentation techniques such as collision-induced dissociation (CID) and electron-transfer/associated dissociation (ETD/EAD) offer structural insights by generating fragment spectra, known as MS2, MS/MS or MSn spectra ^11–13^. While simple full scan MS techniques can yield molecular mass information of glycans, critical for deducing monosaccharide compositions, distinguishing isobaric compounds – molecules sharing identical masses – presents a challenge, which requires additional separation methods to enhance specificity, as these cannot be distinguished by MS1 information alone ^14^. To resolve these ambiguities, techniques like liquid chromatography (LC)^15^ and capillary electrophoresis (CE)^16,17^ are frequently hyphenated with a mass spectrometer via an electrospray ionization (ESI) interface to provide separation based on the unique physicochemical properties of the glycans. Ion mobility, based on the collision cross section (CCS) of molecules, adds an additional layer of separation within the MS itself^15^. Together, these methods improve identification and quantification, while fragment spectra are used to further refine structural assignments. The inclusion of synthetic standards, whether added to the sample or analyzed separately, enhances accuracy, enabling comparisons of retention/migration times (RT/MT), molecular masses, and fragmentation patterns between known and unknown glycans ^18,19^.

Despite the wide range of approaches available, glycan analysis remains challenging due to the complexity of (CE/LC-)MS data sets. Each MS spectrum consists of a dense mixture of overlapping compound signals, with the number of datapoints varying from tens to thousands, which further complicates the data analysis process. Moreover, the quantity of generated spectra can increase rapidly during LC-MS or CE-MS experiments, with outputs ranging from several thousand to tens of millions of spectra. This results in datasets that often encompass several tens to hundreds of gigabytes of raw signal data. Manually analyzing such extensive data sets is cumbersome, time-consuming, and often infeasible, which highlights the need for sophisticated and advanced bioinformatics tools capable of fully automating the glycan identification and quantification process. While full automation has been achieved for the identification and quantification of peptides and proteins ^20^, this level of automation has yet to be realized for glycans.

The complexity associated with glycan analysis arise primarily from their unique challenges when compared to peptides and proteins. These challenges include the overlapping masses of isobaric monosaccharides, the intricately branched architectures of glycan structures, and the myriad types of glycosidic bonds, all of which present additional layers of complexity that current methodologies have not yet fully addressed ^21^. Nevertheless, recent technological advancements enable semi-automated analysis of these molecules. Researchers are now able to generate a list of compound peaks from raw data using software tools such as MZmine^22^, PASTAQ^23^, XCMS^24^, or proprietary software from MS manufacturers. This generated list typically contains key characteristics, including the observed mass and RT/MT of intact glycans and their fragments, which can be used to cross-reference with MS/MS spectral libraries containing highly confident molecular identifications. Despite these advancements, the workflow remains cumbersome. It often necessitates rigorous manual data verification at each step and requires interfacing between various function and modules of software that do not exhibit seamless interoperability. Consequently, researchers frequently need to adjust output and input files, which can hinder the overall analysis efficiency and induce errors.

To address these limitations, an optimal automation solution of the glycan analysis process would effectively integrate various advanced MS data processing and analysis functionalities to enhance both the efficiency and accuracy of glycomics data assignment (see **Fig. 1**). The tool should feature a comprehensive, built-in glycan composition search space creator, enabling users to build combinatorial or custom glycan libraries with a wide range of chemical modifications. Putative glycan signals should be automatically identified within raw MS1 data, creating dedicated extracted ion chromatograms/electropherograms (EIC/EIE) of all detected glycans. It is essential to accurately annotate monoisotopic peaks and charged state of isotope envelopes to reduce the need for manual data curation. Moreover, the process should incorporate a sophisticated deconvolution algorithm that efficiently deisotopes and resolves different adducts. The implementation of automated accurate peak quantification methods, such as calculating the area under the curve (AUC) for identified glycan peaks within the respective EIC/EIE, along with options for automatic normalization based on internal standards, is crucial for enhancing reproducibility. The workflow must also facilitate the simultaneous analysis of multiple samples, thus enabling direct comparisons of sample and sample groups across data sets. Automation to calculate quality criteria metrics is paramount, including features like fitting of isotopic distribution peaks on the detected isotopologues of a glycan, scoring of chromatogram/electropherogram peak shapes, and calculation of mass accuracy errors to provide a thorough quality assessment for user interpretation, thus allowing for the exclusion of poorly quantified and identified peaks. A user-friendly graphical user interface (GUI) would aid in streamlining the process, making these complex tasks accessible for wet laboratory users. Integration with (existing) databases is essential for improving data management and comparison. The outcomes should be exportable in a human-readable format suitable for interpretation and publication, which includes ready-to-publish cartoon figures of glycans that adhere to the symbol nomenclature for glycans (SNFG)^25^. Additionally, automatic annotation of fragment spectra is necessary for enhancing result interpretability, resulting in a holistic platform that drives innovation in glycomics research with minimal user input.

**Figure 1.**
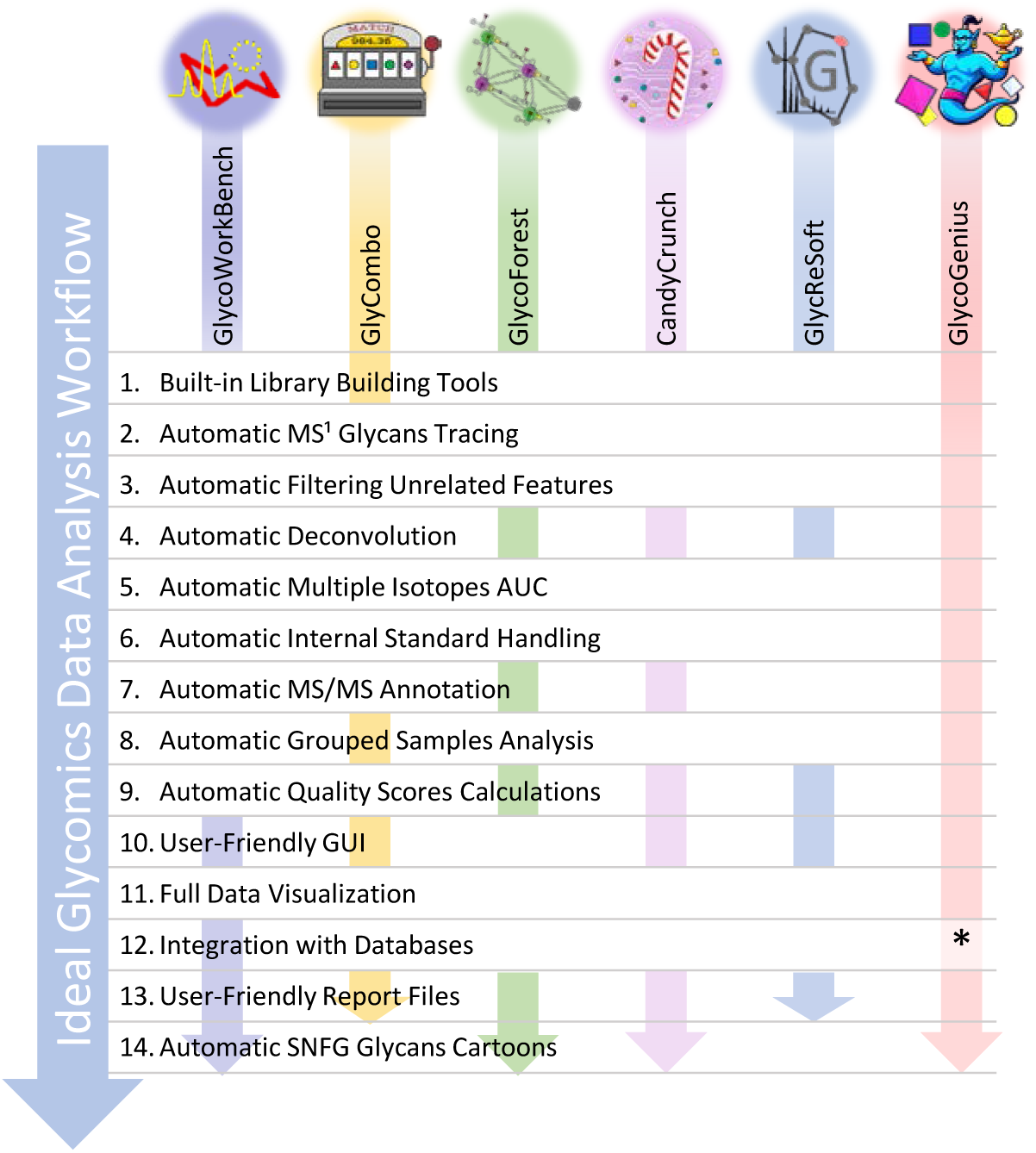
Features and tools required for comprehensive automated glycomics analysis. This scheme shows the most widely used tools and program features required for comprehensive automated analysis of CE/LC-MS glycomics data. *\* This is a desired feature, currently the library and result files generated by GlycoGenius can be easily incorporated into databases to ensure flexible and simple data accessibility*.

Existing tools like CandyCrunch^26^, GlycoForest^27^, GlyCombo^28^, GlycoWorkbench^29^ and GlycReSoft^30^ provide advanced glycomics data analysis functionality but fall short of full automation and integration (**Fig. 1**). For example, GlycoWorkbench is a feature-rich tool, which is able to generate glycan cartoons, explore glycan structure databases, consider diverse chemical modifications and consult whether mass-to-charge ratio (*m/z*) values match known glycan compositions. It is frequently employed for verifying manually analyzed data, but its lack of automation for LC/CE-MS data analysis limits its practical utility. GlycoForest lacks quantification metrics and relies solely on MS/MS data, which may not always be available in standard raw glycomics files, potentially limiting its overall utility. Similarly, CandyCrunch requires MS2 data and utilizes deep learning, exposing the reliability of its identifications on the quality and glycan coverage of the training data set. GlyCombo faces similar challenges, manifested by the absence of data visualization and the limited flexibility of incorporating certain chemical modifications and reducing-end tags for glycans. GlycReSoft, while efficient for single spectra file analysis and offering the most feature-rich functionality among these tools, could as well greatly benefit from enhanced data visualization capabilities. Currently, it only displays EICs, making the data verification process challenging and manual processing work intensive. The tool is unable to automatically integrate findings from different files within a data set, requiring users to manually compile the findings from different samples. Although its quality scoring is robust, it still requires extensive manual verification of the identifications. Additionally, it does not allow for automated quantification of different chromatographic/electropherographic peaks within EICs/EIEs, which is crucial for identifying glycan isomers. It also does not support internal standard quantification or accommodate sialic acid derivatizations. These limitations highlight a notable gap in existing tools, emphasizing the need for a more comprehensive and user-friendly platform for glycomics data analysis, enabling researchers to unlock the full potential of glycomics in understanding the role of glycosylation in health and disease.

In this work, we present GlycoGenius (GG, **Fig. 2**), an advanced program designed to achieve the ideal workflow for comprehensive glycomics LC/CE-MS(/MS) data analysis. The tool integrates feature rich data visualization, encompassing everything from raw spectra and chromatograms/electropherograms to custom-traced EICs/EIEs and isotopic envelope visualization of identified glycans within a unified interface. GlycoGenius employs a streamlined workflow (**Fig. 3**, details available in the *Online Methods* section) that efficiently constructs glycan compositional libraries, filtered EIC/EIE traces, and MS2 annotation. It delivers comprehensive reports and publication-ready figures, enhancing the reliability of results while simplifying data verification for a wide range of glycan classes, such as *N-*glycans, *O-*glycans, glycosaminoglycans (GAGs) as well as glycopeptides datasets within a single easy-to-use user-friendly program compatible with all existing major operational systems such as Windows, Linux and Mac.

**Figure 2.**
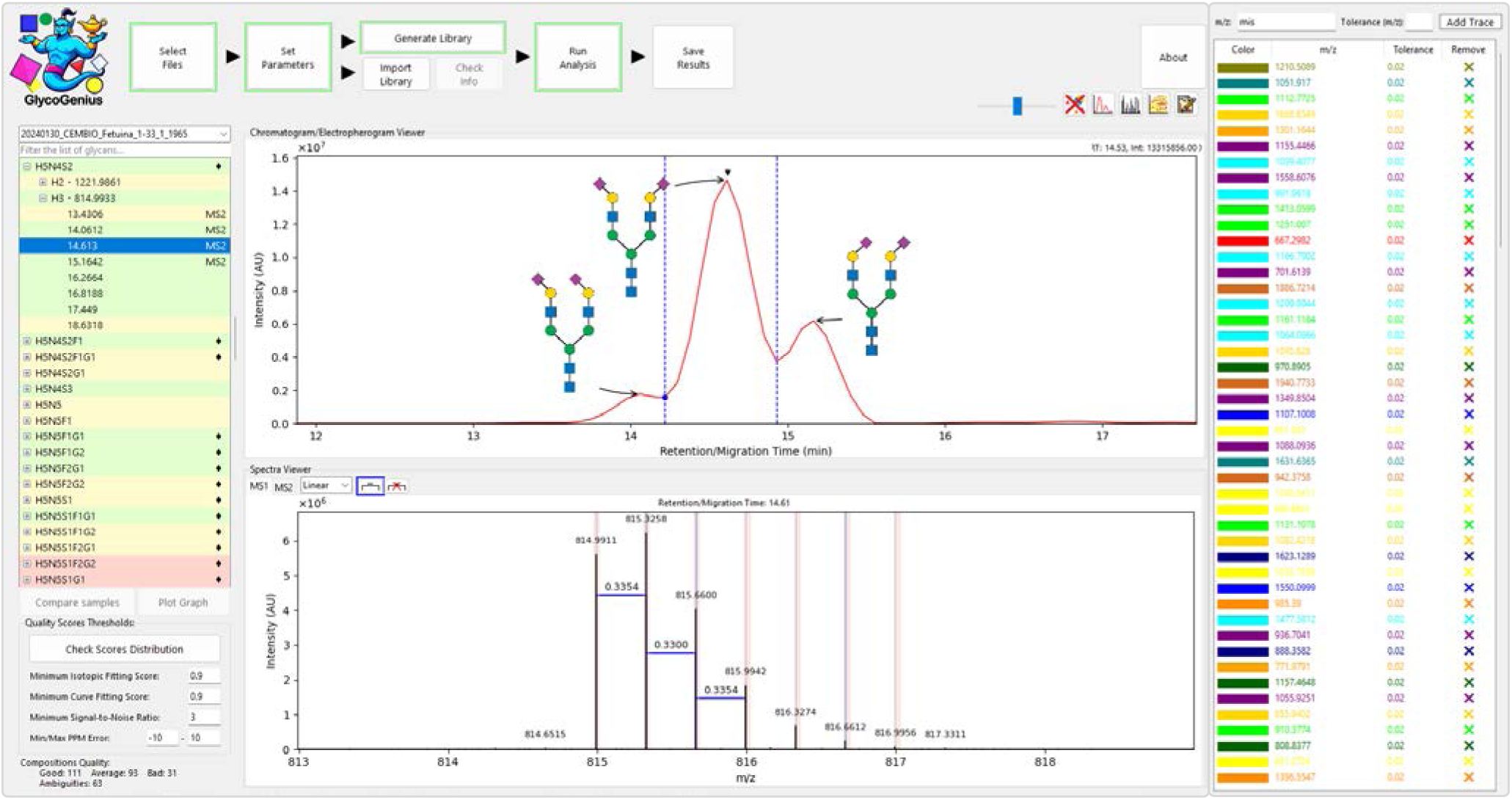
GlycoGenius Graphical User Interface (GUI) Main Window and Quick-Traces. The GUI’s main window features a top menu with step-by-step workflow buttons. Identified glycan compositions are displayed in the left panel; selecting a composition reveals their EIC/EIE on the *Chromatogram/Electropherogram Viewer* (middle panel). Detailed peak information can be accessed by double-clicking on the RT/MT in the left panel. Ambiguous glycans that exhibit the same mass are marked with a black diamond, while peaks with annotated MS2 spectra are marked with a clickable “MS2” label. Selecting a specific RT/MT on the *Chromatogram/Electropherogram Viewer* will show the corresponding spectrum in the *Spectra Viewer* (middle lower panel), highlighting selected isotopic envelope peaks. The right panel features the *Quick Traces* menu for tracing specific *m/z* values and customizing EIC/EIE colors. Users can select multiple glycan compositions or traces by holding the CTRL or SHIFT key.

**Figure 3.**
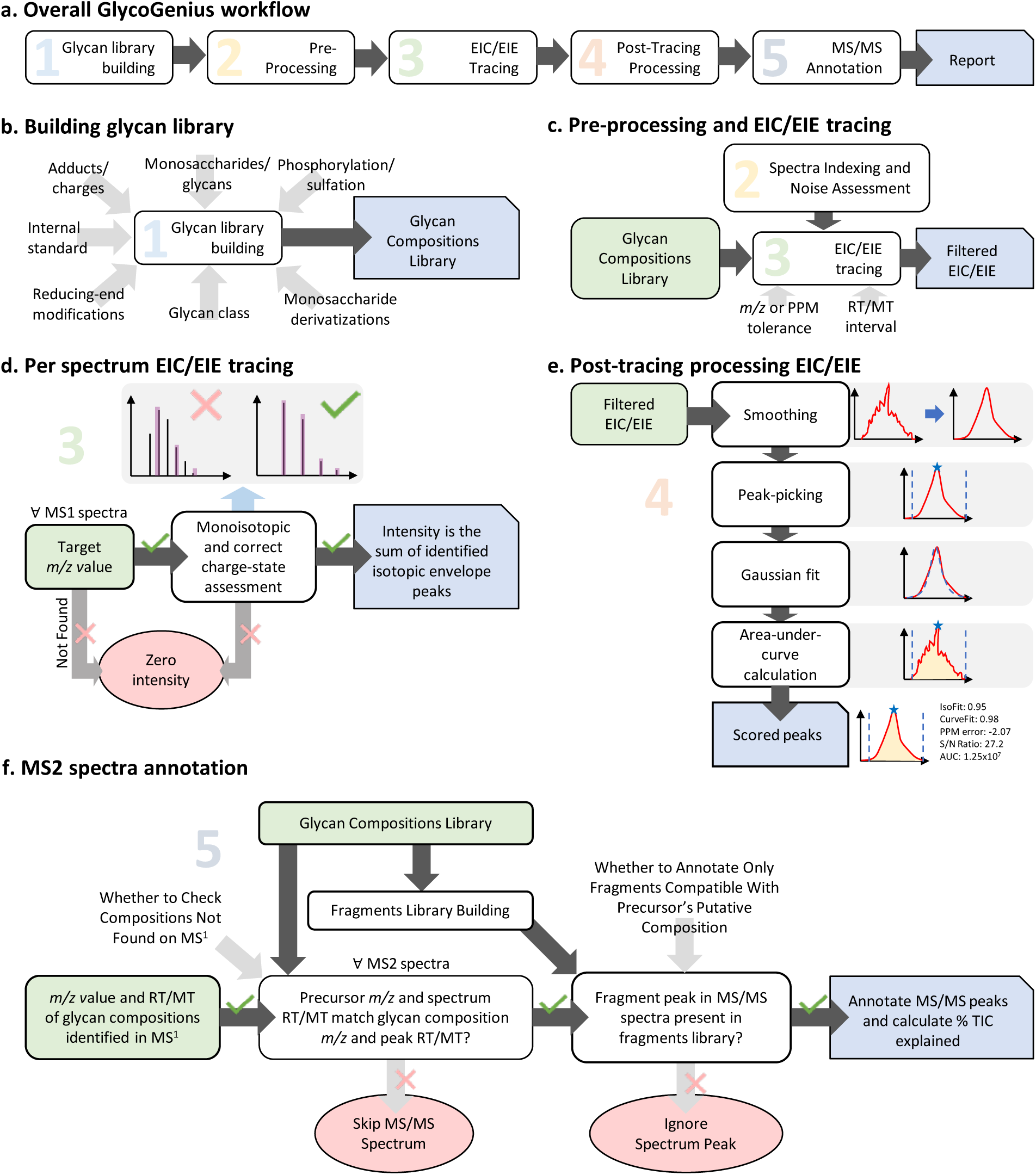
Scheme of GlycoGenius data processing and analysis workflow. **a**, Main modules of the GG analysis pipeline. **b**, Inputs and outputs for building the glycan composition library. **c**, Pre-processing and tracing EICs/EIEs from the glycan library. **d**, Detailed analysis of EIC/EIE traces on an individual MS1 spectrum. **e**, Post-tracing processing of EIC/EIE along with score assessment. **f**, Comprehensive annotation of MS/MS spectra. *Unboxed text: user input; rounded-square boxed text: automated processes; chipped-square blue boxed text: module output. Green boxes indicate input that is automatically pipelined from a previous step output. Red ovals reflect poor outcomes of a given module, which is removed*.

## RESULTS

Two datasets covering different glycan types and analytical conditions were selected to demonstrate the performance, features, and versatility of GG. The first data set encompasses the total plasma *N-* glycome (TPNG) from Lageveen-Kammeijer and de Haan, et al. published in 2019 ^31^. In this study, *N*-glycans were enzymatically released from plasma samples using PNGase-F, followed by sialic acid derivatization through amidation and ethyl-esterification, alongside the modification of the reducing end with a permanent cationic tag (Girard’s reagent P; GirP). The derivatized samples were then analyzed using a CE device, which was coupled with a quadrupole time-of-flight (QTOF) mass spectrometer operating in positive ionization mode. The second data set includes released *O*-glycans derived from keratinocytes, as reported by de Haan, et al. in 2022 ^32^. Here, the glycans were chemically released, enriched using hydrazide beads and tagged with 2-aminobenzoamide (2-AB) at the reducing end. Subsequently, the samples were separated through nanoflow LC on a C18-based analytical column, which was then coupled to an Orbitrap mass spectrometer operated in positive ionization mode. All the glycan compositions shown in this article follow the **GG Monosaccharides Code** (**Table 1**), which uses a simple letter combination (one to two letters) to identify each monosaccharide, followed by the amount of it, with exception of sialic acid modifications.

**Table 1.** GG Monosaccharides Code.

| Monosaccharide | Abbrev. | GG Code | Comment |
| --- | --- | --- | --- |
| Hexose | Hex | H |  |
| Hexosamine | HexN | HN |  |
| *N*-Acetylhexosamine | HexNAc | N |  |
| Deoxyhexose | dHex | F | from Fucose |
| *N*-Acetylneuraminic Acid | Neu5Ac | S | from Sialic Acid |
| *N*-Glycolyl-Neuraminic Acid | Neu5Gc | G |  |
| Uronic Acid | UroA | UA |  |
| Xylose | Xyl | X |  |
| Amidation | Am | Am[G] | α2,3 Sialic Acid |
| Ethyl-Esterification | EE | E[G] | α2,6 Sialic Acid |
| Reducing End (Including Tag) | N/A | T | Only in fragments |

### Total plasma *N-*glycan detection by GlycoGenius

In the original publication, the data underwent manual screening for *N*-glycan compositions documented in the literature^33–38^, encompassing 500 different *N*-glycan compositions. The assignments were based on the exact mass and migration order of these *N*-glycans. Subsequently, a targeted data analysis was performed on aligned raw “mzXML” files using an adapted version of LaCyTools v1.0.1 build 8^39^. The article reported a total of 167 *N-*glycan identifications (158 unique compositions), which included the differentiation of α2,3 and α2,6 linked sialic acids, and 49% of the identified *N*-glycans were confirmed by MS2. Using GG, an automated analysis of the same data (n=3) revealed 174 unique *N*-glycan compositions, all meeting the established quality thresholds (isotopic envelope fitting Score > 0.8, curve fitting score > 0.5, −20 < PPM error < 20, signal-to-noise (S/N) > 3, detailed in the *Data Processing* section found under *Online Methods*). This analysis took one hour and 50 minutes to complete on a computer equipped with a 6-core/12-threads CPU (detailed relevant specifications provided in the *Data Processing* section, under *Online Methods*).

From the 158 *N-*glycan compositions found in the original article, 115 of those were confirmed by GG with the established quality thresholds. The remaining 43 compositions were analyzed in detail and fitted into six categories (**Fig. 4b**). Most compositions fell into four categories (74.4%; 32 compositions) representing glycans detected by GG, but that failed to meet the isotopic fitting quality threshold to varying degrees. Particularly the isotopic envelope fitting scores of these compositions were below 0.8 or detection was inconsistent across replicates (a requirement imposed to replicate the original publication analysis conditions). Another category included glycan compositions that could be found manually, but were missed by GG (16.2%; 7 glycan compositions). The remaining compositions were not found during the manual search on the raw data (9%; 4 glycan compositions). Manual search of the seven *N-*glycans not detected by GG revealed that they were of low abundance (relative abundance < 0.09%, as reported in the original publication) and had very specific particularities that made their signals hard to identify reliably even on a manual inspection. One of these *N*-glycans was identified in the original publication associated with an unknown adduct, of which proper molecular formula and matching mass could not be replicated in this study or using other glycan analysis platforms; one was mislabeled in the original publication (correctly identified by GG as H8N3E1 instead of H8N3Am1, data not shown); three had inconsistent signals with evidence on the presence of the glycan in three or fewer spectra, barely forming an electropherographic peak; two had their isotopic envelope heavily mixed with another unknown isotopic envelope. Interestingly, GG identified 59 *N-*glycan compositions that were not identified in the original publication (**Fig. 4a**). Among these, 46 compositions were absent from the literature that was explored in the original publication ^33–38^. A complete list of the identified compositions on each data set can be found in **Supplementary Table 1.**

**Figure 4.**
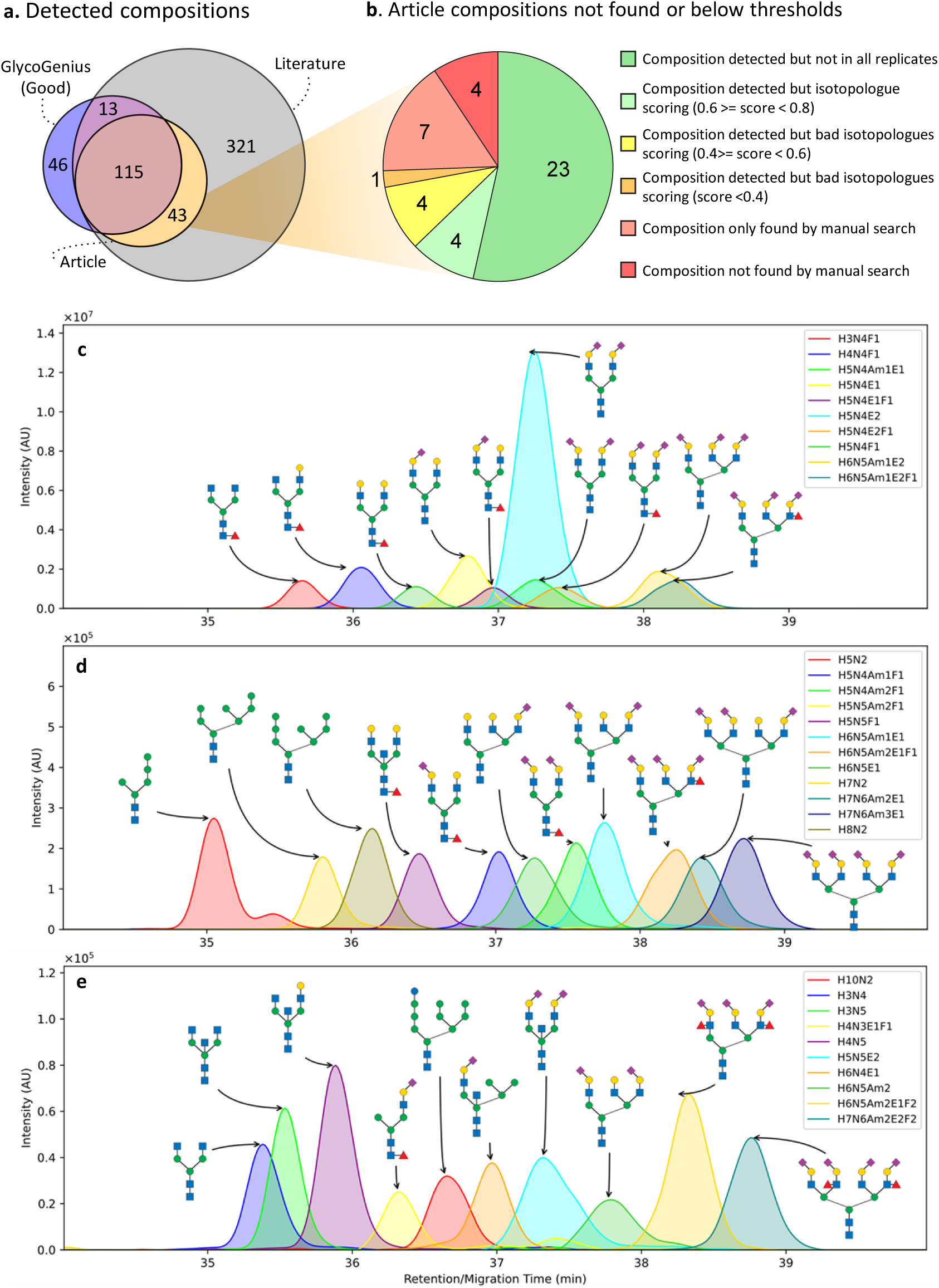
Glycan structure annotation of total plasma CE-MS/MS *N*-glycan profile obtained by manual identification and GlycoGenius. **a,** Venn diagram illustrating the overlap of unique *N*-glycan compositions found in literature, those detected manually in the original article, and the *N*-glycans identified automatically by GG. **b,** *N*-glycan compositions not detected or below GG quality criteria were examined manually and classified into six detection categories. **c,** EIE of the 10 most abundant *N*-glycans. **d,** *N*-glycans with an abundance between 0.5% to 1%. **e,** *N-*glycans under an abundance of 0.25%. Putative glycan structures detected by GG displayed on **c**, **d** and **e**.

Using the *Draw* module, GG is able to automatically generate cartoons of the glycan structures in the SNFG format in the *Chromatogram/Electropherogram Viewer*. These figures replicated the findings of the original study, including the 10 most abundant *N*-glycans (**Fig. 4c**, *Fig. 2d* from Lageveen-Kammeijer and de Haan et al., 2019). Additionally, figures for the EIEs of 10 *N*-glycans with relative abundance and between 0.5% and 1% (**Fig. 4d**) and below 0.25% (**Fig. 4e**) were also generated.

Overall, GG successfully replicated the results found by manual data evaluation performed by glycomics experts. It detected 151 *N-*glycan compositions from the 158 published in the original article, with 115 considered of good quality. Notably, GG identified 59 additional *N-*glycan compositions not found in the original publication, leading to a total of 174 compositions meeting all quality criteria thresholds. From these 59 additional compositions, 46 that were not documented in the explored literature.

### Keratinocytes *O-*glycan detection by GlycoGenius

In the original publication, data analysis began with feature detection using the Minora Feature Detector node in Thermo Proteome Discoverer 2.2.0.388 (Thermo Fisher Scientific). The generated peak list was exported from Thermo Proteome Discoverer and imported to GlycoWorkbench 2.1 (build 146), where it was matched to glycan compositions ranging from 0 to 8 hexoses, 0 to 8 *N*-acetylhexosamines, 0 to 3 fucoses, 0 to 4 *N*-acetylneuraminic acids, and the 2-AB reducing end label. Further matching included additional compositions containing 0 to 6 hexoses, 0 to 6 *N*-acetylhexosamines, 0 to 2 fucoses, 0 to 2 *N*-acetylneuraminic acids, 0 to 3 pentoses and the 2-AB label. The list of identified compounds was then imported to Skyline^40^ via the *Molecule Interface*, facilitating the generation of EICs for the first three isotopologues of each glycan composition. Chromatographic peaks were selected manually based on criteria including accurate mass (within a −1 PPM to 1 PPM error range), isotopic dot product (> 0.85), and minimal intensity threshold (1×10^6^), with compositions required to meet these parameters in only one of the samples (n=19). The original study ultimately identified 27 unique *O*-glycan compositions. A complete list of the identified *O*-glycan compositions on each dataset can be found in **Supplementary Table 2**.

The automated analysis performed by GG identified 25 *O*-glycan compositions that met the predefined quality criteria (isotopic envelope fitting score > 0.8, curve fitting score > 0.5, −20 < PPM error < 20, S/N > 3, details can be found in the *Data Processing Section*, under *Online Methods*). The analysis was completed in approximately one hour and 40 minutes on a computer equipped with a 6-core/12-threads CPU (detailed relevant specifications are provided in the *Data Processing Section*, under *Online Methods*). In comparison, an automated analysis using GlycReSoft revealed a total of 39 compositions within the quality threshold (score > 8) recommended by the software developers ^41^. The preprocessing (a step within the software that requires user input for each sample file) took roughly 30 min, and the identification of glycans (another step within GlycReSoft that requires user input for each sample file) spanned two hours and 30 minutes, resulting in a total analysis time of approximately three hours on the same computer, and this estimate still excludes the time spent browsing menus and the time required to write scripts to convert the glycan list to the necessary GlycReSoft import format and to merge the results from multiple samples into a single glycan quantitative table.

Initial comparisons between the article analysis, GG and GlycReSoft analyses (**Fig. 5a-c**) revealed that, from the 27 reported *O*-glycan compositions in the original publication, GG identified 16 meeting the quality criteria (**Fig. 5b**). The remaining 11 *O*-glycans were successfully detected by GG, but failed the quality criteria regarding isotopic fitting score to various extents (**Fig. 5c**), suggesting that these compounds may not possess the atomic compositions corresponding to the putative glycans. Additionally, GG identified 9 *O*-glycan compositions that were not reported in the original article, reaching a total of 25 identifications.

**Figure 5.**
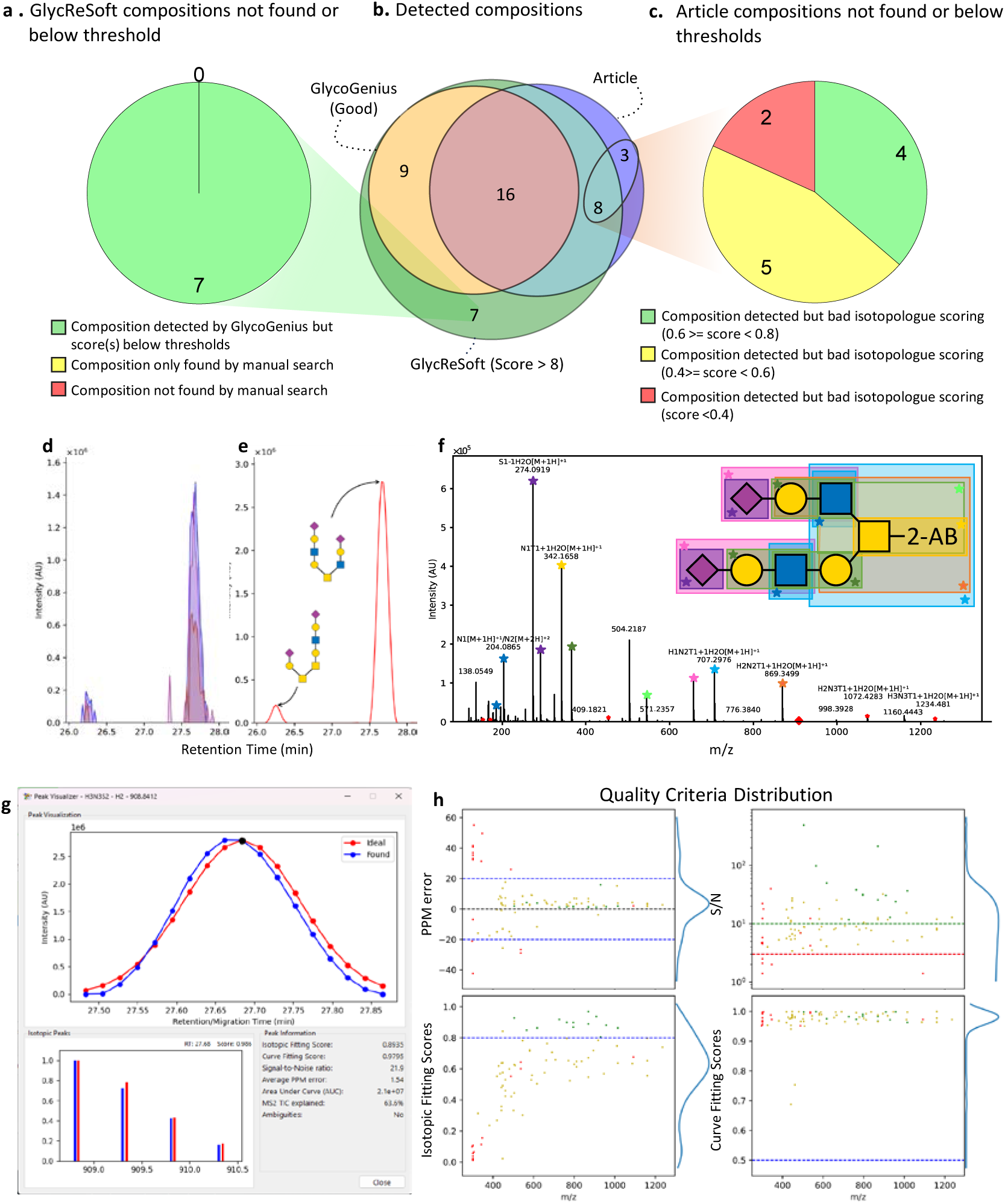
Glycan structure annotation of LC-MS/MS *O*-glycan profile obtained by GlycoGenius and GlycReSoft. **b,** Venn diagram illustrating the overlap of *O*-glycans manually identified in the article, automatically identified by GG and/or by GlycReSoft. **a,** In-depth examination of 7 *O*-glycan compositions identified by GlycReSoft, but not found by GG or because of scores below set thresholds. **c,** Detailed analysis of 11 *O*-glycans compositions identified in the original article but not found by GG or with scores below thresholds. **d,** EIC of three isotopologues of the *O*-glycan with composition H3N3S2 [M+2H]^+2^ (*m/z* 908.8412), traced by GG using the *Quick-Trace feature* and a 0.02 *m/z* threshold. **e,** EIC of H3N3S2 automatically traced by the GG. **f,** Annotated fragment spectrum of the second peak of H3N3S2. Peaks marked with a star indicate automated annotation by GG. Specific fragments are marked with color-coded stars, corresponding to the highlighted fragments in the manually added glycan cartoon on the top (for more details see **Extended Data Table 1**). **g**, *Peak Visualizer* for the second peak of H3N3S2, showing curve fitting scoring (top), the isotopic envelope fitting scoring at the maxima of the peak (bottom, left) and the qualitative and quantitative attributes of the peak (bottom, right). **h,** Visualization of quality criteria distribution of a representative sample of the dataset, peak quality thresholds are indicated by colored dots; green dots fulfill all quality thresholds, yellow dots failed one quality threshold and red dots failed two or more quality thresholds.

GlycReSoft identified a total of 39 *O*-glycan compositions, with 23 aligning to those reported in the original article (**Fig. 5b**), and successfully recognized all *O*-glycans identified by GG. Additionally, GlycReSoft detected 7 *O*-glycan compositions that met the recommended score threshold of the tool, but were not found with satisfying quality scores by GG nor in the original publication. A detailed investigation revealed that while GG had detected these compositions, they did not pass the isotopic fitting score criterion (**Fig. 5a**). To illustrate the filtering performed by GG on the EIC tracing, the composition H3N3S2 was analyzed using the *Quick-Trace* feature (**Fig. 5d**). This trace is comparable to the EIC found in the original publication, which was done using Skyline^40^ (*Fig. 5a* of the original publication). The filtered EIC automatically traced by GG (**Fig. 5e**) is compared to the *Quick-Trace*, showcasing deconvoluted, smoothed and filtered peaks, as made evident by the absence of the thin signal observed at approximately 27.4 minutes in **Fig. 5d**, which only contains the second isotopic peak of the isotopic envelope. Fragmentation of the most abundant peak allowed GG to automatically annotate approximately 64% of the Total Ion Current (TIC) of the MS2 spectrum (**Fig. 5f, Extended Data Table 1**). Users can access detailed characteristics of a peak through the *Peak Visualizer* by double-clicking the identified chromatographic peak in the glycan list on the left side of the window (**Fig. 5g**).

**Extended Data Table 1.**
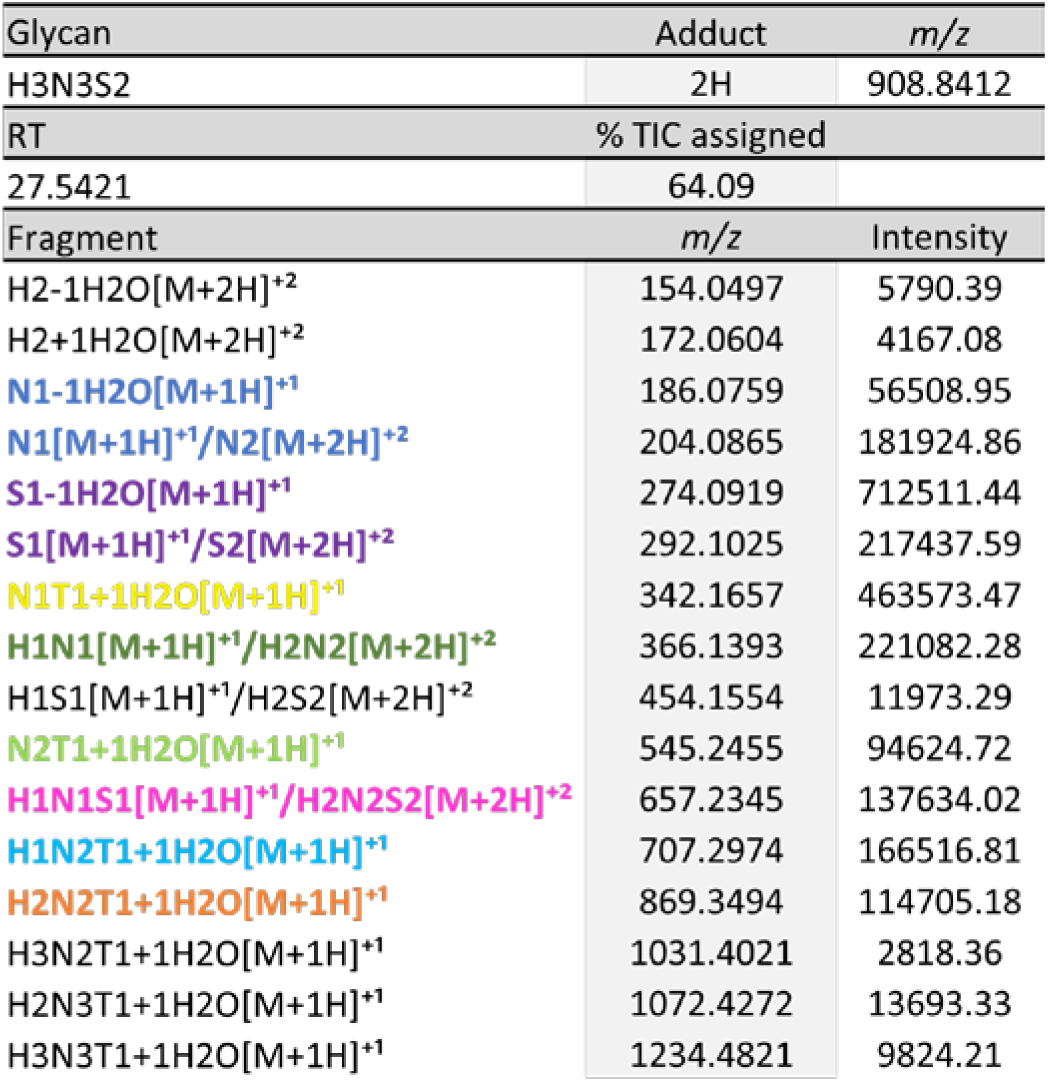
Automatically Annotated H3N3S2 Fragments.

Overall, GG successfully identified all *O*-glycan compositions reported in the original article and those recognized by GlycReSoft. The quality scoring standards automatically filtered many low confidence assignments based on their isotopic envelope distribution, as shown by the representative quality criteria distribution, which is accessible from the GUI (**Fig. 5h**). The analysis ultimately resulted in a total of 25 *O*-glycan composition identifications within the quality criteria stipulated, all of which were manually verified.

## DISCUSSION

Here, we introduce GlycoGenius, a new bioinformatic tool designed for the analysis of LC/CE-MS(/MS) glycomics data. This innovative platform provides an end-to-end solution for glycan identification and quantification, capable of processing multi-sample data sets, within a single program. GlycoGenius streamlines the entire workflow, from library construction to final results reported in user-friendly formats that includes human-readable tables of quantitative and qualitative information, PDF report files, and high-resolution figures. The tool enables not only replication of previous findings, but also the identification of novel glycan compositions with unparalleled efficiency and accuracy, while filtering out dubious identifications (**Fig. 4a**). Using the built-in functionalities of GG, we effortlessly replicated figures from previous publications using newly acquired data sets. In particular, the *GG Draw* module simplifies the creation of engaging visuals, enabling users to swiftly generate high-quality graphics. It also facilitates the automatic generation of SNFG-compliant cartoons, providing clear and accurate representations of the putative glycan structures with minimal effort.

From the analyzed data sets, GG successfully detected most *N-*glycans and all the *O-*glycans compositions reported in the original publications. Its scoring system deemed most of the detections to be high-quality identifications. Additionally, for the *N*-glycome study, GG identified 59 *N-*glycans compositions not found in the original publication, with 46 of them absent from the explored literature. The seven *N-*glycans missed by GG, they were of very low relative abundance, with two of them detected in the original publication under specific conditions that could not be replicated using GG library building, GlycoWorkbench or manually (unknown adduct and unmatching mass). Other missed glycans exhibited severely compromised signal, such as heavily disturbed isotopic envelope or inconsistent signal throughout the MT of the eluting peak.

Notably, one glycan, which had been misidentified in the original publication (as H8N3Am1), was correctly identified as H8N3E1 by GG. This identification matches the mass reported in the original article, and meets all quality thresholds (data not shown). One of the glycan EIE displayed in the original publication (*Figure 2f*, yellow electropherogram), is reported as a glycan of composition H5N5E2, as opposed to the correct identification of H5N5Am2. Four of the reported identified *N*-glycans were not found within the data, even after a manual inspection of the raw data. These misidentifications highlight the necessity of automated glycan composition assignments as human errors are prone to happen and can be effectively avoided through automating LC-MS(/MS) data processing by GG.

GlycoGenius and GlycReSoft are among the few available tools capable of addressing key challenges in glycomics data analysis, such as automatically generating a search space for glycans, detecting them in MS1-only datasets, generating EICs/EIEs, quantifying the abundance of compositions, and implementing a scoring system for meticulous examination of the data quality. The primary distinction between the analysis conducted by GG and GlycReSoft, aside from the shortened analysis time by GG, lies in the level of data quality scrutiny. While the developers of GlycReSoft recommended a score of 8 or higher for identifying glycans, this threshold is insufficient to reliably determine whether an isotopic envelope can be attributed to a glycan or to a different compound with same mass. This limitation comes from the fact that the scoring system used by GlycReSoft consists of four different metrics, of which the individual score ranges and their impact on the data are not clearly defined. As a result, two compounds may achieve a total score equal to or higher than 8 while having widely different individual scores, reducing the confidence of these identifications and requiring thorough manual verification of the results. In contrast, GG uses mainly the isotopic fitting score to determine the identity of a compound. A score of 0.8 or higher is considered a high-confidence identification, provided the compound is within the expected PPM error range, but this value can also be adjusted to meet particular needs in the analysis. Conversely, if this scoring criterion is not met, the monoisotopic *m/z* value and charge state may align with the target molecule, yet they could still represent an isobaric compound unrelated to the glycan. Furthermore, GlycReSoft does not detect different chromatographic/electropherographic peaks within each EIC/EIE, making it difficult to identify isomers. It also requires users to employ additional scripting or manual data management to merge the outcome of multiple samples into one large quantitative glycan composition table. Depending on the data requirements (i.e.: what information, such as scores, RT/MT, AUC, etc., from the results the user wants within their unified dataset) and programming expertise of the user, this process can take from tens of minutes to several hours of extra effort. Moreover, the LC-MS(/MS) data pre-processing and glycan identification phases in GlycReSoft take significantly more time for multiple samples when compared to GG. For example, in the *O-*glycan data set, GlycReSoft required 3 hours to finish the analysis whereas GG completed the same task in 1 hour and 40 minutes. Additionally, data assessment at spectrum level requires the use of external software to access the raw data and manually search for spectra and other features that corresponds to the reported identifications, further increasing the overall workload of the user, whereas GG supports native spectrum, isotopic envelop and EIC/EIE visualization.

GlycoGenius effectively addresses the challenges commonly encountered in glycomics data analysis, offering innovative solutions to streamline workflows and enhance research outcomes. It facilitates generating quantitative glycan tables that can be loaded in MetaboAnalyst^42^, allowing users e.g. to detect differential levels of glycans in different sample groups and perform other statistics such as hierarchical clustering of samples with similar glycan compositions. This integration allows for advanced data visualization through heatmaps, volcano plots, principal component analysis (PCA), receiver operating characteristic (ROC) curves, and more. Notably, this interoperability is unique among glycomics data analysis software tools, providing researchers with a powerful bioinformatic tool to expedite the analysis of larger datasets. The cutting-edge algorithms employed by GG simplify the handling of extensive data sets, making it possible for researchers of all levels of expertise to obtain meaningful insights. This application not only enhances the accuracy of glycan identification, but also offers robust relative quantification capabilities essential for comparative studies. Furthermore, its interactive visualization features enable researchers to thoroughly examine trends in glycosylation patterns such as glycome changes associated with pathological diseases. By breaking down the barriers to advanced glycomics data analysis, GG empowers scientists to delve into the intricacies of glycosylation and its roles in health and disease. This empowerment has the potential to significantly accelerate glycan profile related discoveries, enhancing our understanding of disease mechanisms and identifying prospective therapeutic targets. By enhancing both accuracy and efficiency, GG represents a significant step forward in unlocking the full potential of glycomics for biomedical research.

While GG currently provides robust and comprehensive data analysis capabilities, additional features and improvements are in development for future implementation. Although glycoforms of a single glycopeptide can be efficiently analyzed by GG, full glycoproteomics-level analysis has not yet been achieved. Integrating GlycReSoft^30^ glycopeptide identification results as an input spectral library is planned and would allow for glycoproteomics data analysis using all the power of currently implemented GG functionalities. Additionally, GG currently lacks the ability to automatically determine the precise structure for a given glycan composition which is desired for generating the glycan cartoons for publication. To address this, integration of MS2 data annotation with structure-elucidating powers of CandyCrunch’s^26^ into GG is being studied. Furthermore, the implementation of the following features is planned: spectra file RT/MT trimmer; an enhanced RT/MT alignment tool powered by PASTAQ^23^; and integration with glycan structure databases, such as GlyTouCan^43^. These upgrades will not only expand the analysis capabilities of GG but also improve the user experience and data processing accuracy, further paving the way of GG as a leading tool in glycomics research.

## ONLINE METHODS

### Datasets

The total plasma *N*-glycome dataset, originally published by Lageveen-Kammeijer and de Haan et al., 2019^31^, is available on MassIVE (identifier MSV000083478)^44^. The keratinocyte *O*-glycome data set, published in de Haan N. et al., 2022^32^, was downloaded from the PRoteomics IDEntifications Database (PRIDE, identifier PXD029644)^45^.

### Data Processing

For the total plasma *N*-glycome dataset, the following parameters were used for library building in GG: Monosaccharides: 5 to 22; Hexoses: 3 to 10; HexNAcs: 2 to 8; Sialic acids: 0 to 4; Deoxyhexoses: 0 to 2; *N*-acetyl neuraminic acids: 0 to 4. Other monosaccharides were set to zero. The *Force Glycan Class* option was set to “*N-glycans*”. The generated library was combined with the library containing 500 *N*-glycans compositions used in the original publication and any duplicates were removed, leading to a total of 2070 unique compositions (1769 unique masses). The reducing end tag was set to “*GirP*” and the “*Amidated/Ethyl-Esterified*” sialic acids derivatization option was toggled. Hydrogen (proton) was the only adducts chosen, with charges ranging from 1 to 3 per molecule. Other library settings were left at default values (see the online manual at the GitHub repository, available under the *Code Availability* section).

For the analysis settings (accessible via the right panel of the *Parameters window*, in the GUI), the tolerance unit chosen was *m/z* value, and the maximum tolerance was set to 0.02 *m/z.* The samples were analyzed with a MT range between 35 and 50 minutes. Default settings were maintained for the remaining parameters. The quality criteria thresholds used for glycan composition identification were: isotopic fitting score: 0.8; curve fitting score: 0.5; signal-to-noise ratio (S/N): 3; parts-per-million (PPM) error: from −20 to 20, and only *N*-glycans found within all three replicates were considered. The rest of the settings were left at the default values.

The keratinocyte *O-*Glycans data set library was built using the following parameters in GG: Monosaccharides: 1 to 10; Hexoses: 0 to 8; HexNAcs: 0 to 8; Sialic acids: 0 to 4; Deoxyhexoses: 0 to 2; *N*-acetyl neuraminic acids: 0 to 4, and Xyloses: 0 to 3. Other monosaccharides were set to zero. The *Force Glycan Class* option was set to “*O-glycans*”. To ensure that the library covered all the *O*-glycans found in the original article, it was supplemented with the *O*-glycan compositions identified in there and any duplicates were removed, resulting in a library containing 247 unique *O*-glycan compositions (and masses). The reducing-end tag chosen was “*2-AB*” and hydrogen (proton) was the only adduct selected, with charges ranging from 1 to 3 per molecule. The remaining library settings were left at the default values.

The analysis settings included a maximum mass tolerance of *m/z* 0.02 and the analyzed RT interval was from 20 to 35 minutes. All other settings were left at default values. A second run was performed for a single *O*-glycan composition of H3N3S2 using identical settings, with the “*Analyze MS2*” and “*Only assign fragments compatible with the precursor composition*” options turned on and “*Look for fragments of glycans not found on MS1*” option left off. These settings ensured that GG initially validated the presence of the glycan in the MS1 spectra, verified the correct isotope distribution, and identified the MS2 spectra of which the precursor *m/z* matched H3N3S2. MS2 peaks were annotated only if their fragment putative compositions were compatible with the precursor putative composition.

All analyses used the “*Multithreaded*” option with number of CPU Cores set to “*all*”, which uses all of the available CPU threads minus 2 (i.e.: a CPU with 12 threads will use 10 threads if CPU Cores is set to “*all*”). The analyses were performed on a computer with the following specifications: CPU: AMD Ryzen 5 3600 @ Stock speeds (6-core, 12-threads); RAM: 32 GB DDR4-3000; Storage: Samsung 980 PRO 1TB SSD.

### Design and Functionality of GlycoGenius

GlycoGenius is entirely developed in Python (version >= 3.6) and uses multiple packages to deliver its robust functionality: Pyteomics^46,47^ (used for accessing raw data files and performing precise mass calculations of the mass and isotopic distributions of atomic formulas), Dill^48,49^ (facilitates pickling, which is the conversion of Python objects into byte streams, allowing Python objects to be saved to files that can be reloaded by GG on demand), Numpy^50^ (used for mathematical calculations within the workflow), SciPy^51^ (used in the Whittaker-Eilers’ smoothing^52^ and in the alignment algorithms), and Pandas^53,54^ (used for exporting the results to Excel files).

GlycoGenius features a versatile interface designed for a wide range of users. Its GUI is ideal for performing data analysis and visualization for users with no or limited Python and bioinformatics knowledge, providing an intuitive platform for data analysis and visualization (**Fig. 2**). Additionally, a command-line interface (CLI), useful for performing analysis on headless remote computers and to batch process data sets using custom scripts, is available. This flexibility allows GG to run on any modern computation clusters and cloud system, thus enabling the analysis of large datasets on high-performance computing infrastructure.

Every peak picked from EICs/EIEs during the analysis can be examined in detail using an integrated peak viewer. This feature displays comprehensive information, including per spectrum isotopic distribution fitting scores, overall isotopic distribution scores, peak curve fitting visualization and score for S/N, AUC, PPM error, and percentage of TIC explained, if MS2 spectra were annotated for the peak in question (**Fig. 5g**). Users can also visualize the score distributions of all parameters within the GUI. By adjusting score thresholds for specific parameters, researchers can dynamically filter and refine the data (**Fig. 5h**). Moreover, the user can visually inspect each EIC/EIE and the associated MS1 and MS2 spectra in the GUI, allowing for in-depth characterization and evaluation.

The analysis is saved in the “*gg*” format (GlycoGenius file format), which preserves all the information from the analysis, including chromatograms, scores, detected peaks of the isotopic envelope, curve and isotopic fittings, and MS2 annotations. When loaded together with the source spectra file(s) (i.e.: the “*mzML*” or “*mzXML*” file(s) used in the analysis), these files also allow visualization of the spectra directly within GG. The “*gg*” files can be uploaded to raw, meta and processed data sharing platforms, such as GlycoPOST^55^, Zenodo^56^, MassIVE^57^, or PRIDE^45^. This compatibility enables seamless sharing and validation of analysis results, fostering collaboration and reproducibility in glycomics research.

### Description of GlycoGenius Analytical Workflow

The GG workflow (**Fig. 3a**) is designed to facilitate the identification and analysis of glycan compositions in LC/CE-MS(/MS) data sets. The process begins with the creation of a glycan composition library. Users can provide monosaccharide ranges or custom lists of compositions, derivatizations, and adducts/charges ranges. The software calculates the *m/z* values and theoretical isotopic distributions based on the natural abundance of atom isotopes (**Fig. 3a**, 1). The spectra data files, in “*mzML*” or “*mzXML*” standard formats developed by Human Proteome Organization Proteomics Standards Initiative (HUPO PSI), are loaded into the program, and the software indexes the spectra and estimates noise-levels (**Fig. 3a**, 2). Using the generated glycan composition library, GG explores the spectra and traces EICs/EIEs, assessing how well the MS1 data follows the glycan structure isotope distribution (**Fig. 3a**, 3). The EICs/EIEs are smoothed, and peaks are detected based on local maxima and minima and scored for isotopic distribution fitting against the theoretical distribution. The peaks detected in the EICs/EIEs are fitted against Gaussian curves, and the S/N is calculated based on the monoisotopic peak of the spectrum at the maximum of the chromatographic/electropherographic peak. The AUC is calculated based on the unsmoothed data to retain precision (**Fig. 3a**, 4). Then, the glycan composition library *m/*z values are checked one by one against the MS2 spectra precursor *m/z* values, and if a match is found within a chosen tolerance window, the MS2 spectrum is analyzed peak by peak against a library of fragments built from the original glycan composition library. Thus, spectrum peaks matching *m/z* values of fragments are appropriately annotated (**Fig. 3a**, 5).

### Glycan Composition Library Generation Algorithm

The glycan composition library is generated by GG based on user-defined inputs that customizes the library for specific data analysis needs. The required inputs include: a range for the monosaccharides or a list of glycans using the GG Monosaccharides Code (**Table 1**); the mass, chemical composition, or simplified GG Monosaccharides Code of an internal standard, if one is used; the range of adducts and maximum number of charges per molecule; the reducing-end modifications that will be applied to the glycans (including the internal standard, if a glycan is entered); other monosaccharide derivatizations, such as permethylation or sialic acid differentiation; range of phosphorylation and/or sulfation per glycan; and the glycan class constraint, if desired (**Fig. 3b**).

If a range of monosaccharides is specified, mathematical combinatorics will be applied using the monosaccharides to create all the possible combinations of compositions. When a glycan class constraint (*N*-glycans, *O-*glycans or GAGs) is specified, the compositions are limited by the rules specified in **Extended Data Table 2**. Alternatively, a list of glycans can be manually input by the user or imported from a comma-or line-separated text file (.txt). In this case, only the glycan compositions on the list will be generated.

**Extended Data Table 2.** Glycan Classes Monosaccharides Constraints.

| Category | Hexoses<br>(H) | N-acetyl<br>hexosamines<br>(N) | Sialic acids<br>(S+G) | Deoxyhexoses<br>(F) | Pentoses<br>(X) | Hexosamines<br>(HN) | Hexuronic<br>acids (UA) | Additional<br>constraints |
| --- | --- | --- | --- | --- | --- | --- | --- | --- |
| N-Glycans | $\geq 2$ | $\geq 2$ | $\leq 2 \times (N - 2)$<br>and $\leq H - 2$ | $< N$ | $\leq 1$ | 0 | 0 | |
| O-Glycans | $\leq N + 1$ | $\leq H + 3$ | $\leq H + N$ | $\leq H + N$ | 0 | 0 | 0 | |
| GAGs | $H + UA \leq N + HN$ | $HN + N \leq UA + H + 1$ | 0 | 0 | 0 | $HN + N \leq UA + H + 1$ | $H + UA \leq N + HN$ | Both H and UA > 0 not allowed |

Reducing-end tag modifications are applied to all generated glycan compositions. Options include a reduced end, a reducing-end tag (such as 2-aminobenzamide or 2-AB, 2-aminobenzoic acid or 2-AA, procainamide or ProA, GirP, etc.) selected from a list or with a specified added mass or molecular formula, or no modification. The user can also choose to apply permethylation or sialic acids derivatization (currently only amidation/ethyl-esterification are supported) to the generated glycan compositions. Ranges for phosphorylation and/or sulfation can be specified, and the glycan compositions will be generated with all the possibilities within the specified ranges (i.e.: for the glycan H5N4S2 and phosphorylation ranging from 0 to 1 and sulfation from 1 to 2, the compositions H5N4S2+1(s), H5N4S2+1(s)+1(p), H5N4S2+2(s), H5N4S2+2(s)+1(p) is generated).

Once all the glycan compositions and their modifications and adducts are defined, GG calculates the theoretical isotopic envelope distribution based on two available options: 1) slow mode, which provides precise results by taking the available isotopes of all the atoms (carbon, oxygen, nitrogen, hydrogen, etc.) into consideration, but requires considerable computation time, depending on glycans molecular sizes, or 2) fast mode, which considers only the carbon isotopes and applies empirically derived correction equations (**Supplementary Table 3**) to approximate the results of the slow option. This significantly reduces computation time, while maintaining accuracy for most applications.

Sulfation and phosphorylation are considered in the isotopic envelope calculation, regardless of whether the slow or fast option is used. If the slow isotopic envelope distribution is calculated, a ‘*high-resolution’* option can be chosen, which allows consideration of minor effects in the isotopic distribution, such as the small difference in the mass added to the monoisotopic peak when replacing an ^16^O isotope with one ^18^O (≈ 2.004245 Da) and replacing 2 ^13^C with two ^12^C ones (≈ 2.003242 Da), which can only be detected in ultra-high-resolution MS.

The built library is saved in “*ggl*” file format (GlycoGenius library file), which can be imported later to use in subsequent analyses. Additionally, GG automatically creates a file compatible with Skyline^40^ transitions’ list format, enabling seamless integration with Skyline^40^ workflows. This flexibility ensures that users can efficiently build, customize, and apply glycan composition libraries for diverse data analysis tasks.

### Pre-Processing Algorithm

Before analyzing a sample file using GG, the raw data files must be converted from the vendor format to the “*mzML*” or “*mzXML*” file format and profile data should be centroided. This conversion can be done using the analysis software provided by the specific vendor of the MS device or by using MSConvert from the ProteoWizard toolset^58^. Once the file is loaded into GG, it will be fully scanned to index MS1 and MS2 spectra. This indexing step optimizes multiple functionalities by allowing to selectively analyze either MS1 or MS2 spectra without having to check the MS level of each spectrum during processing. Noise levels are inferred during this step, based on the average of the 66.8^th^ percentile (or two standard deviations) of the non-zero signal intensities from each spectrum (**Fig. 3c**, upper box).

To increase the precision of noise estimation, the noise level of the first and last quarter of the *m/z* range of each spectrum is inferred using the same method and is considered as the noise levels at the beginning and end of the *m/z* range. Throughout the pipeline, the local *m/z* value-specific noise levels are calculated based on a linear interpolation and regression of the noise levels calculated at the beginning and end of the relevant spectrum. This approach ensures precise noise estimation across the full *m/z* range of each spectrum, enabling accurate S/N calculations for every identified glycan.

### Extracted Ion Chromatograms/Electropherograms Tracing Algorithm

For each glycan composition and adduct combination in the library, GG calculates an associated *m/z* value. With this information, GG searches every MS1 spectrum for a peak corresponding to that *m/z*, within a tolerance chosen by the user. If, in a given spectrum, the corresponding *m/z* value is found, GG verifies whether the peak is monoisotopic and its isotopic envelope has the correct charge state. Upon successful verification, the isotopic envelope is checked and scored and the sum of the detected isotopic envelope peaks intensities (up to the theoretical proportion for that glycan) is used as the intensity for that specific RT/MT in the EIC/EIE. If a peak corresponding to the *m/z* value is not found, or the isotopic distribution fails the monoisotopic peak and correct charge state tests, the corresponding RT/MT intensity in the EIC/EIE is set to zero (**Fig. 3d**).

To fail the monoisotopic check, a spectrum peak must fulfill two criteria. The first criterion is that another spectrum peak must exist with an *m/z* value offset negatively by the mass of the hydrogen atom divided by an integer representing the possible charge states (i.e., −1.0074, −0.5037, −0.3328, etc.). For the second criterion, the intensity of this other peak (from first criterion) must exceed a given threshold (i.e., if the peak is small enough, it will be ignored). For the second criterion, the intensity ratio between the first two peaks in the postulated isotopic envelope is evaluated. For this purpose, the average mass and the theoretical isotopic envelope of peptides, glycans and lipids with an increasing number of building blocks were calculated (similar to a simplified Averagine^59^ model, used for peptides, but our model consider also different molecule classes). A linear interpolation and regression were performed to establish the relationship between the mass of these compounds and the relative intensity of the second isotopic envelope peak (**Supplementary Table 4**). Using **Equation 1**, GG evaluates whether two given peaks are the monoisotopic and first isotopologue peaks of the same isotopic envelope.

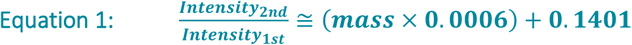

To verify whether the isotopic envelope has the correct charge state, GG examines for the existence of a peak with *m/z* offset positively from the monoisotopic peak by the mass of the hydrogen atom divided by an integer representing the possible charge states. If such a peak is found, it is checked whether its intensity is high enough for it to qualify as the second peak of the isotopic distribution using **Equation 1**. To fail a charge state check, the tested peak must meet three criteria: the positively offset peak must be found; it must be of high enough intensity; and must match a charge state other than the putative glycan charge state.

If the picked *m/z* value is confirmed to be monoisotopic and with the correct charge state, the PPM error of the monoisotopic peak is calculated and the quality of the remaining isotopologues is assessed. The ratio between the theoretical intensity and the measured intensity of each isotopic envelope peak is scaled using a logistic function (**Equation 2**) with the optimized steepness parameter (*k* value) corresponding to 10 and the midpoint set to 0.5.

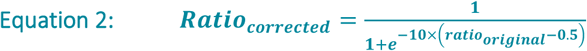

This scaling follows common practice^60^ in scoring and the *k* value of 10 amplifies values over the midpoint and attenuates the values below the midpoint, while keeping a linear range around 0.5. This scaling sets apart low values from high values, creating a clear distinction while still leaving average values open for interpretation. Once the ratios of all the isotopic peaks are calculated and scaled, they are assigned a weighted score using an exponential decay function with a fine-tuned decay factor of 1.25 (**Equation 3**).

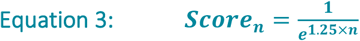

Where *n* is the isotopic peak number (i.e., 1 is the first isotopologue, 2 is the second isotopologue, etc.). This equation generates values that are normalized to result in a sum of 1, and the ratios are used to scale the scores obtained from **Equation 2**.

This adjustment puts a higher weight on the first two isotopic envelope peaks after the monoisotopic peak (71.4% and 20.5%, respectively, thus aggregating 91.9% of the total score on the second and third isotopic envelope peaks). This allows for consistent scoring, as these two peaks are the most intense and stable peaks of the isotopic distribution for glycans, making them more reliable.

### Post-Tracing Processing Algorithm

Once the filtered EIC/EIE is traced, GG applies the Whittaker-Eilers smoothing algorithm^61^. Chromatographic/electropherographic peaks are picked based on local maxima and their boundaries are determined by adjacent local minima or where the intensity reaches a value below 0.01% of the maximum intensity in the EIC/EIE. A Gaussian curve is fitted into the peaks and the coefficient of determination (R²) is used to determine the curve fitting score. With the boundaries of each peak determined, the AUC is calculated using the unsmoothed EIC/EIE. An overall PPM error and isotopic envelope fitting score is obtained with a weighted average procedure, where the scores of each spectrum within the peak boundary is weighted by a transformed version of the Gaussian curve used during the curve fitting scoring. The procedure provides a peak list, which includes peak abundance reflected by AUC of the peak, peak boundaries, isotopic envelope fitting scores, curve fitting scores, PPM error and S/N (**Fig. 3e**).

The Whittaker-Eilers smoother used, as implemented in Python by Midelet J. *et al.*, 2018^52^, is a computationally fast algorithm for smoothing based on a penalized least squares method and with automatic interpolation. Its smoothing capabilities are determined by two factors: the roughness penalty and the order of smoothing. The order of smoothing was optimized to the value of 2 and the roughness penalty (λ) is adjusted dynamically based on the number of data points per minute (dpm) in the chromatogram/electropherogram, using **Equation 4**.

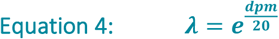

These settings maintain the unique peak shapes of the data, while providing sufficient smoothness for peaks to be effectively picked, without generating bias, as a Gaussian smoothing filter would (i.e., a Gaussian smoothing filter causes all peaks to be Gaussian-shaped, so when fitting a Gaussian against this peak, it will always tend to be a perfect fit).

The Gaussian fitting curve is calculated using the probability density function (PDF) of a Gaussian distribution (**Equation 5**) that is later scaled to the peak height.

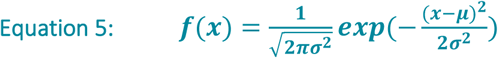

The best fit is found by an iterative process with 10 iterations of the standard deviation (σ) and all possible integer iterations of the mean (μ). The resulting curve is fitted to the peak and the coefficient of determination (R²) is used as the curve fitting score. Its transformed (squared) shape is used as a weight to calculate the PPM error and the isotopic envelope fitting score of the peak. This transformation assures that the most reliable point of the peak (the maximum) has a significantly greater impact on the score of the peak than the rest of the points.

### MS2 Spectrum Annotation Algorithm

For each glycan composition and adduct combination in the library, GG scans the MS2 spectra in the spectra data file to identify potential matches. An MS2 spectrum is selected for annotation if its precursor *m/z* value that matches, within a tolerance set by the user, up to the third theoretical isotopic envelope *m/z* value of the glycan composition and adduct combination. Additionally, if the glycan was identified in MS1, the MS2 spectrum RT/MT must be within the RT/MT boundaries of the peak identified in MS1 in order for it to be selected for annotation. For the annotation, GG generates a library of all the possible fragments based on the glycan composition library, and each peak in the selected MS2 spectrum is checked against this library. If a peak matches an *m/z* value within mass error tolerance of a fragment in the library and the fragment composition is compatible with the glycan composition of the putative precursor glycan, the MS2 peak is annotated (**Fig. 3f**).

The user can choose to only look for glycans in MS2 spectra if they were found on the MS1 first, and to only annotate peaks that are compatible with the putative fragmented glycan composition. The fragment library also includes labeling or reducing chemical modifications of the reducing-end part of the glycan (referred as “*T*” in the GG Monosaccharides Code, **Table 1**). Once the annotation is completed, the percentage of the TIC annotated on that spectrum is calculated and the annotated MS2 spectrum can be viewed in the GUI (**Fig. 5f**), while all the annotated fragments, their *m/z* values and their intensity are available in an Excel table, as shown in **Extended Data Table 1**.

### Additional Features

GlycoGenius offers a variety of features designed to enhance the data analysis and visualization of glycomics data, while providing functionalities for intuitive data exploration and generation of publication-quality outputs. GlycoGenius automatically detects ambiguities in the glycan library, notifying the user if different glycans share the exact same mass. It is also capable of automatically determining the class of *N*-glycans and their ratio (i.e., the percentage of glycans that are *Paucimannose*, *Hybrid*, *Complex* or *High-Mannose*) based on the number of hexoses (H) and *N*-acetyl hexosamines (N). If N is equal to 2 and H is equal to or smaller than 3, the *N-*glycan is classified as a *Paucimannose*; If it is not classified as a *Paucimannose*, and H is greater than N plus one, and N is greater than 2, the *N-*glycan is classified as *Hybrid*; If it is neither a *Paucimannose* nor a *Hybrid*, and N is equal to 2 and H is greater than 3, then the *N-*glycan is classified as *High-Mannose*; If the *N-*glycan does not fit in any of these classes, it will be classified as *Complex*.

To facilitate data exploration, GG provides multiple visualization tools, including a fully interactive two-dimensional map of the LC-MS(/MS) data that plots the RT/MT on the x-axis and the *m/z* values on the y-axis, with the intensities highlighted in a color scale and location of MS/MS spectra acquired in DDA mode (**Extended Data Fig. 1a**). Another alternative data visualization method is showing a composite spectrum called a maximum intensity spectrum (MIS), which is built from the maximum intensity that each *m/z* value achieves through the whole chromatographic/electropherographic run. With this visualization method, the researcher can quickly check for the presence of glycans in the sample and/or for the quality of the library built, as glycans can be identified on this spectrum using a previously built library, and it includes proper isotopic envelope scoring (**Extended Data Fig. 1b**). The user also has the option to trace any *m/z* value using the *Quick-Trace* menu. This feature allows the user to trace specific *m/z* values, with customizable colors, and to plot multiple traces simultaneously by holding CTRL or SHIFT and selecting multiple traces. Multiple *m/z* values can be added in a single trace using semicolons in the *m/z* value field (**Fig. 2**, right side panel).

**Extended Data Figure 1.**
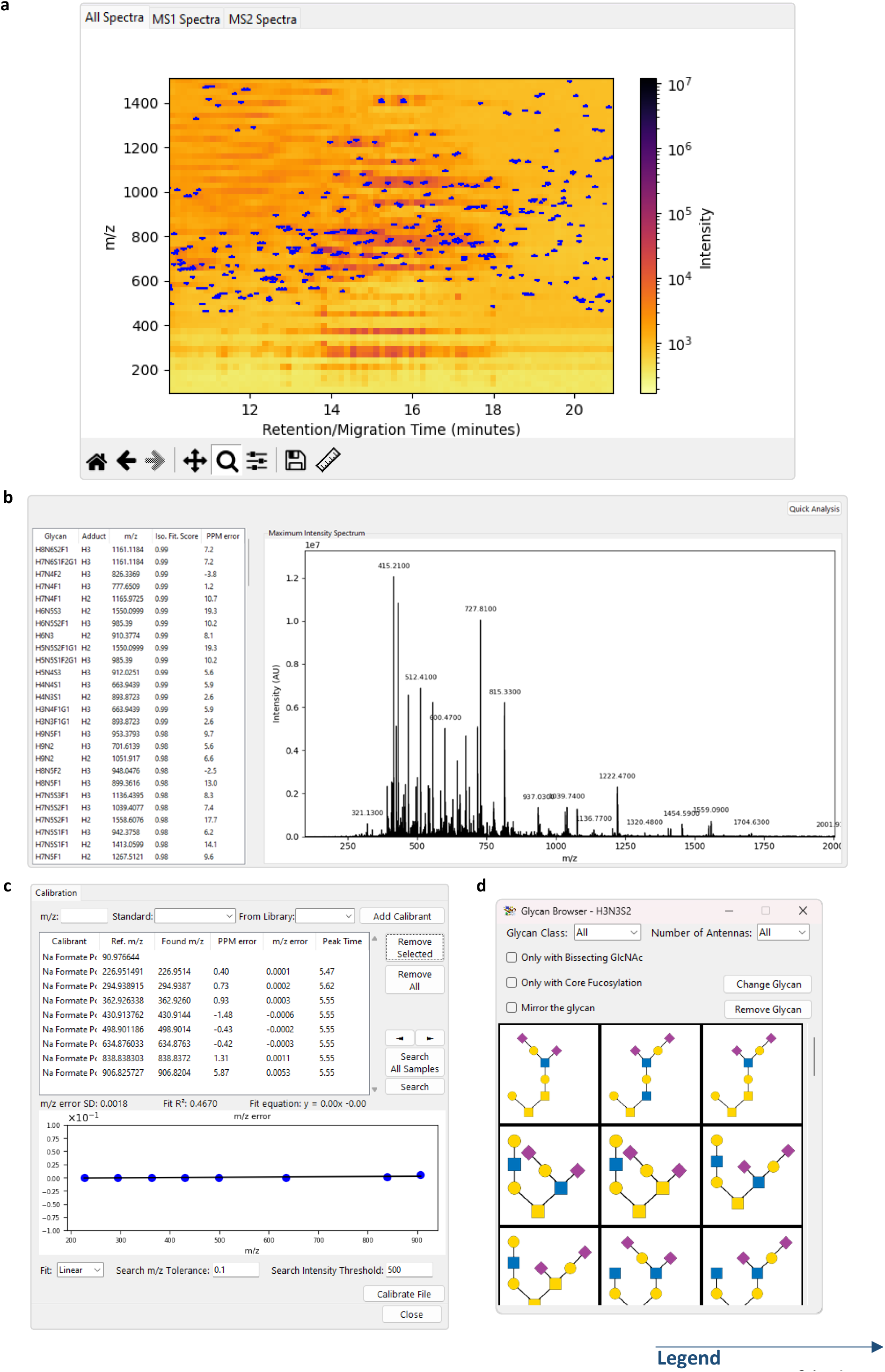
The data visualizations modules in GlycoGenius, File Editor and GlycoGenius Draw. **a**, Two-dimensional heat map, where the x-axis represents RT/MT and the y-axis the m/z values. Peak intensity is displayed in a color scale of the heatmap cell, from dark red (most intense) to white (less intense), in logarithmic scale. Blue markings indicate peaks chosen for fragmentation in MS1 spectra. **b**, Maximum Intensity Spectrum (MIS), which is a composite spectrum constructed from the maximum MS1 intensity of m/z values in the LC-/CE-MS. A quick analysis can be performed on this spectrum to quickly assess the presence of glycan compositions, depicted on the left side panel. **c**, GlycoGenius File Editor, depicting the calibration module. A list of calibrants is composed by the user based on the top options (m/z, standard or from glycans library) and searched within the sample. After adjusting the hits, the file is calibrated, generating a report. **d**, The Glycan Browser window allows the user to filter out choices and select the glycan cartoon that best matches the putative glycan or remove it altogether.

GlycoGenius features a MS file editor that currently only supports spectrum *m/z* calibration (**Extended Data Fig. 1c**), with RT/MT trimming (remove unwanted RT/MT ranges and save to file only desired parts) and a retention time alignment based on PASTAQ^23^ implementation of the Warp2D^62^ algorithm planned in the future. In the calibration module, the user can input custom *m/z* values, calibrant standards, or select glycans from their library to create a list of internal calibrants, which will be searched for in the whole spectra file. Once the search is done, every found calibrant will display the theoretical mass, the detected mass, the mass error (in absolute *m/z* values and PPM), and the RT/MT at which the compound was found at highest intensity. Users can refine the *m/z* calibrant list and perform either quadratic or linear calibration, generating a new calibrated “mzML” file and a detailed PDF report on the calibration details.

Glycan cartoons following the SNFG^25^ guidelines can be drawn automatically by GG using its *Draw* module (**Fig. 2**, visible in the chromatogram/electropherogram viewer). This module includes precalculated antenna combinations of *N-* and *O*-glycans based on known biosynthetic pathways. This allows for quick local generation of the glycan images to be used by GG. When the *Draw* module is activated, glycans will generate their cartoons when selected and display them on the *Chromatogram/Electropherogram Viewer*. The cartoons can be freely rearranged on the canvas by dragging them, allowing users to optimize the layout. Additionally, double-clicking on a cartoon enables users to modify it by selecting a different structure for the same glycan composition or removing the cartoon entirely (**Extended Data Fig. 1d**). This functionality is specifically designed to facilitate the creation of publication-quality figures, ensuring that the visual representation meets high scientific and aesthetic standards. The automatically generated cartoons for the glycan compositions may require user adjustments to reflect the correct putative structure.

## DATA AVAILABILITY

All datasets converted to mzML format and GG results files generated during data analysis and presented in this work are available on Zenodo (https://doi.org/10.5281/zenodo.14630509).

## CODE AVAILABILITY

GlycoGenius v. 1.2.8 (with GUI v. 1.0.7) was used in this study and is available on GitHub at https://github.com/LoponteHF/GlycoGenius_GUI, under GNU GPL v3+ open-source license.

## ACKNOWLEDGEMENTS

This work was supported by funding of Coordenação de Aperfeiçoamento de Pessoal de Nível Superior (CAPES), awarded to H.F.L. and the Dutch Research Council (NWO) that funded the X-omics Road Map program (project 184.034.019) and project VI.Veni.222.262, awarded to G.L.K. The authors are grateful for the opportunities and support afforded by the Sector Plan Pharmaceutical Sciences, which was implemented in the overarching Sector Plan Beta II, put into action by the Dutch Ministry of Education, Culture and Science (OCW). The authors thank the Centro de Espectrometria de Massas de Biomoléculas (CEMBIO, UFRJ, Rio de Janeiro, Brazil).

## AUTHOR INFORMATION

These authors contributed equally: Adriane R. Todeschini, Peter L. Horvatovich and Guinevere S. M. Lageveen-Kammeijer.

### Contributions

H.F.L. performed literature research on the topic, designed the software, wrote the program code, analyzed the datasets, drew the figures, wrote and revised the manuscript and user-guide and manages the GitHub repository; J.Z. tested the program extensively, participated actively in glycomics scientific discussion, participated in the program user experience (UX) design, revised the user-guide and provided test datasets for the development of GG; Y.D. participated in the coding design of the *GG draw* module; I.A.O. tested the program, participated extensively in glycomics scientific discussion and the program UX design and provided test datasets for the development of GG; K.B. set up the test server for GG at Analytical Biochemistry, provided IT infrastructure support and UX feedback; A.R.T. participated in the initial idealization of the GlycoGenius development, providing a name and logo idea, participated in scientific discussion regarding glycobiology, provided UX design feedback, and revised the user-guide; P.L.H. and G.S.M.L.K. participated actively and extensively in the software design, from algorithm to UX design, in scientific discussion regarding glycomics and data analysis, provided test datasets, drew and revised the figures and revised the user-guide. All authors revised the manuscript.

## ETHICS DECLARATIONS

### Competing interests

Authors declare no competing interests.

## Notes

https://doi.org/10.5281/zenodo.14630509

https://github.com/LoponteHF/GlycoGenius_GUI

### Competing Interest Statement

The authors have declared no competing interest.

