## Supplementary Tables for "GlycoGenius: the ultimate high-throughput glycan composition identification tool"

### Table of Content Supplementary Tables

**Supplementary Table S-1:**

Identified *N* -glycan composition by original publication, literature used in orginal publication and GlycoGenius.

**Supplementary Table S-2:**

Identified O-glycan composition by original publication (Art. Id), GlycReSoft software tool (GlycReSoft Ids) and GlycoGenius (GG Ids).

**Supplementary Table S-3:**

Corrections calculated for fast isotopic distribution calculation.

**Supplementary Table S-4:**

Model to estimate intensity of second isotopic envelope peak of a given molecule.

| Supplemenatry Table S-1. Identified *N* -glycan composition by original publication (Art. Id), literature used in orginal publication (Lit.Ids) and GlycoGenius (GG Ids). | | | |
|---|---|---|---|
| Glycan | Art. Ids | GG Ids | Lit. Ids |
| H10N2 | X | X | X |
| H2N3F1 | X | X | X |
| H3N2 | X | X | X |
| H3N3 | X | X | X |
| H3N3E1 | X | X | X |
| H3N3F1 | X | X | X |
| H3N4 | X | X | X |
| H3N4F1 | X | X | X |
| H3N5 | X | X | X |
| H3N5F1 | X | X | X |
| H4N2 | X | X | X |
| H4N3 | X | X | X |
| H4N3Am1 | X | X | X |
| H4N3Am1F1 | X | X | X |
| H4N3E1 | X | X | X |
| H4N3E1F1 | X | X | X |
| H4N3F1 | X | X | X |
| H4N4 | X | X | X |
| H4N4Am1 | X | X | X |
| H4N4Am1F1 | X | X | X |
| H4N4E1 | X | X | X |
| H4N4E1F1 | X | X | X |
| H4N4F1 | X | X | X |
| H4N5 | X | X | X |
| H4N5E1 | X | X | X |
| H4N5E1F1 | X | X | X |
| H4N5F1 | X | X | X |
| H5N2 | X | X | X |
| H5N3 | X | X | X |
| H5N3Am1 | X | X | X |
| H5N3Am1F1 | X | X | X |
| H5N3E1 | X | X | X |
| H5N3F1 | X | X | X |
| H5N4 | X | X | X |
| H5N4Am1 | X | X | X |
| H5N4Am1E1/H5N7 | X | X | X |
| H5N4Am1E1F1/H5N7F1 | X | X | X |
| H5N4Am1E1F2/H5N7F2 | X | X | X |
| H5N4Am1F1 | X | X | X |
| H5N4Am2 | X | X | X |
| H5N4Am2F1 | X | X | X |
| H5N4E1 | X | X | X |
| H5N4E1F1 | X | X | X |
| H5N4E2 | X | X | X |
| H5N4E2F1 | X | X | X |
| H5N4F1 | X | X | X |

| | | | |
|---|---|---|---|
| H5N5 | X | X | X |
| H5N5Am1E1F1/H5N8F1 | X | X | X |
| H5N5E1 | X | X | X |
| H5N5E1F1 | X | X | X |
| H5N5E2 | X | X | X |
| H5N5E2F1 | X | X | X |
| H5N5F1 | X | X | X |
| H6N2 | X | X | X |
| H6N3 | X | X | X |
| H6N3Am1 | X | X | X |
| H6N3E1 | X | X | X |
| H6N3F1 | X | X | X |
| H6N4Am1 | X | X | X |
| H6N4Am1E1/H6N7 | X | X | X |
| H6N4E1 | X | X | X |
| H6N5Am1E1/H6N8 | X | X | X |
| H6N5Am1E1F1/H6N8F1 | X | X | X |
| H6N5Am1E2/H6N8E1 | X | X | X |
| H6N5Am1E2F1/H6N8E1F1 | X | X | X |
| H6N5Am1E2F2/H6N8E1F2 | X | X | X |
| H6N5Am1F1 | X | X | X |
| H6N5Am2 | X | X | X |
| H6N5Am2E1/H6N8Am1 | X | X | X |
| H6N5Am2E1F1/H6N8Am1F1 | X | X | X |
| H6N5Am2E1F2/H6N8Am1F2 | X | X | X |
| H6N5Am2F1 | X | X | X |
| H6N5Am3 | X | X | X |
| H6N5Am3F1 | X | X | X |
| H6N5E1 | X | X | X |
| H6N5E1F1 | X | X | X |
| H6N5E2 | X | X | X |
| H6N5E2F1 | X | X | X |
| H6N5E3 | X | X | X |
| H6N5E3F1 | X | X | X |
| H7N2 | X | X | X |
| H7N6Am1E1 | X | X | X |
| H7N6Am1E1F1 | X | X | X |
| H7N6Am1E2 | X | X | X |
| H7N6Am1E2F1 | X | X | X |
| H7N6Am1E3 | X | X | X |
| H7N6Am1E3F1 | X | X | X |
| H7N6Am2 | X | X | X |
| H7N6Am2E1 | X | X | X |
| H7N6Am2E1F1 | X | X | X |
| H7N6Am2E2 | X | X | X |
| H7N6Am2E2F1 | X | X | X |
| H7N6Am2E2F2 | X | X | X |
| H7N6Am2F1 | X | X | X |
| H7N6Am3 | X | X | X |
| H7N6Am3E1 | X | X | X |

| | | | |
|---|---|---|---|
| H7N6Am3E1F1 | X | X | X |
| H7N6Am3E1F2 | X | X | X |
| H7N6Am3E1F3 | X | X | X |
| H7N6Am3F1 | X | X | X |
| H7N6Am4F1 | X | X | X |
| H7N6E1 | X | X | X |
| H7N6E1F1 | X | X | X |
| H7N6E2 | X | X | X |
| H8N2 | X | X | X |
| H8N3E2 | X | X | X |
| H8N7Am1E2 | X | X | X |
| H8N7Am1E3 | X | X | X |
| H8N7Am2E1 | X | X | X |
| H8N7Am2E2 | X | X | X |
| H8N7Am2E2F1 | X | X | X |
| H8N7Am2E2F2 | X | X | X |
| H8N7Am3E1 | X | X | X |
| H8N7Am3E1F1 | X | X | X |
| H9N2 | X | X | X |
| H4N4E2 | | X | X |
| H4N6Am1E1/H4N9 | | X | X |
| H5N3E2 | | X | X |
| H5N5E2F3 | | X | X |
| H5N6E1 | | X | X |
| H6N4Am1F1 | | X | X |
| H6N6Am2 | | X | X |
| H6N6Am3 | | X | X |
| H7N5Am2E1F1/H7N8Am1F1 | | X | X |
| H7N6E3F1 | | X | X |
| H7N7Am4 | | X | X |
| H8N3Am2 | | X | X |
| H8N4Am2 | | X | X |
| H3N2F1 | X | | X |
| H3N4Am1 | X | | X |
| H4N4Am1E1/H4N7 | X | | X |
| H4N4Am1E2F1/H4N7E1F1 | X | | X |
| H4N4F2 | X | | X |
| H4N5Am1 | X | | X |
| H4N5Am1F1 | X | | X |
| H4N5F2 | X | | X |
| H5N3E1F1 | X | | X |
| H5N5Am1 | X | | X |
| H5N5Am1E1/H5N8 | X | | X |
| H5N5Am2F1 | X | | X |
| H5N5F2 | X | | X |
| H6N3E1F1 | X | | X |
| H6N4 | X | | X |
| H6N4Am1E1F1/H6N7F1 | X | | X |
| H6N4E1F1 | X | | X |
| H6N4E2 | X | | X |

| | | | |
|---|---|---|---|
| H6N4F1 | X | | X |
| H6N5 | X | | X |
| H6N5Am1E3/H6N8E2 | X | | X |
| H6N5F1 | X | | X |
| H6N6E1F1 | X | | X |
| H6N6E2F1 | X | | X |
| H7N5Am1E1/H7N8 | X | | X |
| H7N6Am1 | X | | X |
| H7N6Am4 | X | | X |
| H7N6Am4F2 | X | | X |
| H7N6E2F1 | X | | X |
| H7N6E3 | X | | X |
| H7N6E4 | X | | X |
| H8N3Am1 | X | | X |
| H8N4Am1E1/H8N7 | X | | X |
| H8N4Am1E3/H8N7E2 | X | | X |
| H8N4Am3E1F1/H8N7Am2F1 | X | | X |
| H8N4E2 | X | | X |
| H8N7Am1E3F1 | X | | X |
| H8N7Am3F2 | X | | X |
| H8N7Am4 | X | | X |
| H8N7Am4F1 | X | | X |
| H8N7E3 | X | | X |
| H9N4E1 | X | | X |
| H9N8Am2E2 | X | | X |
| H10N3Am1E1/H10N6 | | | X |
| H10N5Am1 | | | X |
| H10N5Am1E1/H10N8 | | | X |
| H10N5Am2 | | | X |
| H10N5E1 | | | X |
| H10N5E2 | | | X |
| H10N6Am1E2F5 | | | X |
| H10N6Am2E1F5 | | | X |
| H10N6Am3F5 | | | X |
| H10N6E3F5 | | | X |
| H10N6F3 | | | X |
| H10N7Am1F4 | | | X |
| H10N7E1F4 | | | X |
| H10N9Am2E1 | | | X |
| H10N9Am3E1 | | | X |
| H10N9Am4 | | | X |
| H10N9Am4F1 | | | X |
| H11N10 | | | X |
| H11N2 | | | X |
| H11N4Am1 | | | X |
| H11N4Am1E3F1 | | | X |
| H11N4Am2E2F1 | | | X |
| H11N4Am3E1F1 | | | X |
| H11N4Am4F1 | | | X |
| H11N4E1 | | | X |

| | | | |
|---|---|---|---|
| H11N4E4F1 | | | X |
| H11N5Am1E3F1 | | | X |
| H11N5Am2E2F1 | | | X |
| H11N5Am3E1F1 | | | X |
| H11N5Am4F1 | | | X |
| H11N5E4F1 | | | X |
| H11N7Am1F4 | | | X |
| H11N7E1F4 | | | X |
| H11N9Am1E1F2 | | | X |
| H12N2 | | | X |
| H12N7Am1E3F1 | | | X |
| H12N7Am1F4 | | | X |
| H12N7Am2E2F1 | | | X |
| H12N7Am3E1F1 | | | X |
| H12N7Am4F1 | | | X |
| H12N7E1F4 | | | X |
| H12N7E4F1 | | | X |
| H13N3Am1F1 | | | X |
| H13N3E1F1 | | | X |
| H13N5Am1F3 | | | X |
| H13N5E1F3 | | | X |
| H13N7Am1F2 | | | X |
| H13N7E1F2 | | | X |
| H14N4Am1 | | | X |
| H14N4Am1F1 | | | X |
| H14N4Am1F3 | | | X |
| H14N4E1 | | | X |
| H14N4E1F1 | | | X |
| H14N4E1F3 | | | X |
| H3N3Am1 | | | X |
| H3N3Am1E1F1/H3N6F1 | | | X |
| H3N3Am1F1 | | | X |
| H3N3Am1F2 | | | X |
| H3N3Am2F1 | | | X |
| H3N3E1F1 | | | X |
| H3N3E2F1 | | | X |
| H3N4Am1F1 | | | X |
| H3N4E1 | | | X |
| H3N4E1F1 | | | X |
| H3N4F3 | | | X |
| H3N5Am1F1 | | | X |
| H3N5E1F1 | | | X |
| H3N6 | | | X |
| H3N7F2 | | | X |
| H3N9 | | | X |
| H4N10F3/H4N7Am1E1F3 | | | X |
| H4N3Am1E1/H4N6 | | | X |
| H4N3Am1E1F1/H4N6F1 | | | X |
| H4N3Am1F2 | | | X |
| H4N3Am2 | | | X |

| | | | |
|---|---|---|---|
| H4N3E2 | | | X |
| H4N3F2 | | | X |
| H4N3F3 | | | X |
| H4N4Am1E1F2/H4N7F2 | | | X |
| H4N4Am1F2 | | | X |
| H4N4Am2 | | | X |
| H4N4F3 | | | X |
| H4N5Am1E1/H4N8 | | | X |
| H4N5Am1E1F2/H4N8F2 | | | X |
| H4N5Am1F2 | | | X |
| H4N5Am2 | | | X |
| H4N5Am2F2 | | | X |
| H4N5E1F2 | | | X |
| H4N5E2 | | | X |
| H4N5E2F2 | | | X |
| H4N6E1F1 | | | X |
| H4N7Am3F2 | | | X |
| H4N7E1 | | | X |
| H5N2F1 | | | X |
| H5N2F2 | | | X |
| H5N3Am1E1/H5N6 | | | X |
| H5N3Am2 | | | X |
| H5N4Am1E1F3/H5N7F3 | | | X |
| H5N4Am1E2F1/H5N7E1F1 | | | X |
| H5N4Am1F2 | | | X |
| H5N4Am2E1F1/H5N7Am1F1 | | | X |
| H5N4Am2F2 | | | X |
| H5N4Am2F3 | | | X |
| H5N4E1F2 | | | X |
| H5N4E2F2 | | | X |
| H5N4E2F3 | | | X |
| H5N4F2 | | | X |
| H5N4F3 | | | X |
| H5N5Am1E1F3/H5N8F3 | | | X |
| H5N5Am1F1 | | | X |
| H5N5Am1F2 | | | X |
| H5N5Am2 | | | X |
| H5N5Am2F3 | | | X |
| H5N5E1F2 | | | X |
| H5N6Am1 | | | X |
| H5N6Am1E1F1 | | | X |
| H5N6Am2F1 | | | X |
| H5N6E2F1 | | | X |
| H5N7Am1E1F1 | | | X |
| H5N7Am2F1 | | | X |
| H5N7E2F1 | | | X |
| H5N8Am1E1 | | | X |
| H5N8Am1E1F1 | | | X |
| H6N11/H6N5Am2E2/H6N8Am1E1 | | | X |
| H6N3Am1E1/H6N6 | | | X |

| | | | |
|---|---|---|---|
| H6N3Am1E1F1/H6N6F1 | | | X |
| H6N3Am1F1 | | | X |
| H6N3Am2 | | | X |
| H6N3E2 | | | X |
| H6N4Am1F3 | | | X |
| H6N4Am2 | | | X |
| H6N4Am2F1 | | | X |
| H6N4E1F3 | | | X |
| H6N4E2F1 | | | X |
| H6N5Am1 | | | X |
| H6N5Am1E1F2/H6N8F2 | | | X |
| H6N5Am1E1F4 | | | X |
| H6N5Am1E4F2/H6N8E3F2 | | | X |
| H6N5Am1F2 | | | X |
| H6N5Am2E3F2/H6N8Am1E2F2 | | | X |
| H6N5Am2F2 | | | X |
| H6N5Am2F4 | | | X |
| H6N5Am3E1/H6N8Am2 | | | X |
| H6N5Am3E2F2/H6N8Am2E1F2 | | | X |
| H6N5Am3F2 | | | X |
| H6N5Am4 | | | X |
| H6N5Am4E1F2/H6N8Am3F2 | | | X |
| H6N5Am5F2 | | | X |
| H6N5E1F2 | | | X |
| H6N5E2F2 | | | X |
| H6N5E2F4 | | | X |
| H6N5E3F2 | | | X |
| H6N5E4 | | | X |
| H6N5E5F2 | | | X |
| H6N5F2 | | | X |
| H6N6Am1 | | | X |
| H6N6Am1E1 | | | X |
| H6N6Am1E1F1 | | | X |
| H6N6Am1E2 | | | X |
| H6N6Am1E2F1 | | | X |
| H6N6Am1F1 | | | X |
| H6N6Am1F2 | | | X |
| H6N6Am1F3 | | | X |
| H6N6Am2E1 | | | X |
| H6N6Am2E1F1 | | | X |
| H6N6Am2E1F2 | | | X |
| H6N6Am2F1 | | | X |
| H6N6Am3F1 | | | X |
| H6N6Am3F2 | | | X |
| H6N6E1 | | | X |
| H6N6E1F2 | | | X |
| H6N6E1F3 | | | X |
| H6N6E2 | | | X |
| H6N6E3 | | | X |
| H6N6E3F1 | | | X |

| | | | |
|---|---|---|---|
| H6N6F3 | | | X |
| H6N7Am1E5F2 | | | X |
| H6N7Am2E4F2 | | | X |
| H6N7Am3E3F2 | | | X |
| H6N7Am4E2F2 | | | X |
| H6N7Am5E1F2 | | | X |
| H6N7Am6F2 | | | X |
| H6N7F2652 | | | X |
| H7N2F1 | | | X |
| H7N3Am1 | | | X |
| H7N3Am1E1/H7N6 | | | X |
| H7N3Am1F1 | | | X |
| H7N3Am1F2 | | | X |
| H7N3Am2 | | | X |
| H7N3E1 | | | X |
| H7N3E1F1 | | | X |
| H7N3E1F2 | | | X |
| H7N3E2 | | | X |
| H7N3F1 | | | X |
| H7N3F2 | | | X |
| H7N4 | | | X |
| H7N4Am1 | | | X |
| H7N4Am1E1F3/H7N7F3 | | | X |
| H7N4Am2F3 | | | X |
| H7N4E1 | | | X |
| H7N4E2F3 | | | X |
| H7N4F2 | | | X |
| H7N5Am1 | | | X |
| H7N5Am1E1F1/H7N8F1 | | | X |
| H7N5Am1E1F3/H7N8F3 | | | X |
| H7N5Am1E2F1/H7N8E1F1 | | | X |
| H7N5Am1F1 | | | X |
| H7N5Am2 | | | X |
| H7N5Am2F1 | | | X |
| H7N5Am2F3 | | | X |
| H7N5Am3F1 | | | X |
| H7N5E1 | | | X |
| H7N5E1F1 | | | X |
| H7N5E2 | | | X |
| H7N5E2F1 | | | X |
| H7N5E2F3 | | | X |
| H7N5E3F1 | | | X |
| H7N5F1 | | | X |
| H7N6Am1E1F2 | | | X |
| H7N6Am1E2F2 | | | X |
| H7N6Am1E3F2 | | | X |
| H7N6Am1F1 | | | X |
| H7N6Am1F2 | | | X |
| H7N6Am1F3 | | | X |
| H7N6Am2E1F2 | | | X |

| | | | |
|---|---|---|---|
| H7N6Am2E3F2 | | | X |
| H7N6Am2F2 | | | X |
| H7N6Am3F2 | | | X |
| H7N6Am4F3 | | | X |
| H7N6E1F2 | | | X |
| H7N6E1F3 | | | X |
| H7N6E2F2 | | | X |
| H7N6E3F2 | | | X |
| H7N6E4F1 | | | X |
| H7N6E4F2 | | | X |
| H7N7Am1E2 | | | X |
| H7N7Am1E3 | | | X |
| H7N7Am2E1 | | | X |
| H7N7Am2E2 | | | X |
| H7N7Am3 | | | X |
| H7N7Am3E1 | | | X |
| H7N7E3 | | | X |
| H7N7E4 | | | X |
| H8N3 | | | X |
| H8N3Am1E1/H8N6 | | | X |
| H8N3Am1E1F2/H8N6F2 | | | X |
| H8N3E1 | | | X |
| H8N3F1 | | | X |
| H8N4Am1E1F1/H8N7F1 | | | X |
| H8N4Am1E1F2/H8N7F2 | | | X |
| H8N4Am1E2/H8N7E1 | | | X |
| H8N4Am1E3F1/H8N7E2F1 | | | X |
| H8N4Am1F2 | | | X |
| H8N4Am2E1/H8N7Am1 | | | X |
| H8N4Am2E2/H8N7Am1E1 | | | X |
| H8N4Am2E2F1/H8N7Am1E1F1 | | | X |
| H8N4Am2F2 | | | X |
| H8N4Am3E1/H8N7Am2 | | | X |
| H8N4E1F2 | | | X |
| H8N4E2F2 | | | X |
| H8N5 | | | X |
| H8N5Am1 | | | X |
| H8N5Am1E1F2/H8N8F2 | | | X |
| H8N5Am1F1 | | | X |
| H8N5Am1F2 | | | X |
| H8N5Am1F3 | | | X |
| H8N5Am1F4 | | | X |
| H8N5E1 | | | X |
| H8N5E1F1 | | | X |
| H8N5E1F2 | | | X |
| H8N5E1F3 | | | X |
| H8N5E1F4 | | | X |
| H8N5F2 | | | X |
| H8N6Am1E1F1 | | | X |
| H8N6Am1E1F2 | | | X |

| | | | |
|---|---|---|---|
| H8N6Am1E2F1 | | | X |
| H8N6Am1E2F3 | | | X |
| H8N6Am1F1 | | | X |
| H8N6Am2E1F1 | | | X |
| H8N6Am2E1F3 | | | X |
| H8N6Am2F1 | | | X |
| H8N6Am2F2 | | | X |
| H8N6Am3F1 | | | X |
| H8N6Am3F3 | | | X |
| H8N6E1F1 | | | X |
| H8N6E2F1 | | | X |
| H8N6E2F2 | | | X |
| H8N6E3F1 | | | X |
| H8N6E3F3 | | | X |
| H8N7Am1E2F2 | | | X |
| H8N7Am1E2F3 | | | X |
| H8N7Am2E1F2 | | | X |
| H8N7Am2E1F3 | | | X |
| H8N7Am3F3 | | | X |
| H8N7E3F2 | | | X |
| H8N7E3F3 | | | X |
| H8N7E4 | | | X |
| H8N7F4 | | | X |
| H9N3F2 | | | X |
| H9N4Am1 | | | X |
| H9N4Am1E2/H9N7E1 | | | X |
| H9N4Am1E3F1/H9N7E2F1 | | | X |
| H9N4Am2E1/H9N7Am1 | | | X |
| H9N4Am2E2F1/H9N7Am1E1F1 | | | X |
| H9N4Am3E1F1/H9N7Am2F1 | | | X |
| H9N5Am1E1F1/H9N8F1 | | | X |
| H9N5Am1E2F3/H9N8E1F3 | | | X |
| H9N5Am2E1F3/H9N8Am1F3 | | | X |
| H9N6Am1 | | | X |
| H9N6Am1E1 | | | X |
| H9N6Am1E1F1/H9N9F1 | | | X |
| H9N6Am1F2 | | | X |
| H9N6Am2 | | | X |
| H9N6E1 | | | X |
| H9N6E1F2 | | | X |
| H9N6E2 | | | X |
| H9N7Am1E1F4 | | | X |
| H9N7Am2F4 | | | X |
| H9N7E2F4 | | | X |
| H9N8Am2E1 | | | X |
| H9N8Am2E1F1 | | | X |
| H10N3E2F1 | | X | |
| H10N5F1 | | X | |
| H2N7F3 | | X | |
| H3N5E1F2 | | X | |

| | | |
|---|---|---|
| H3N5E1F3 | | X |
| H3N6Am1F1 | | X |
| H3N7Am1F1 | | X |
| H4N4Am1E1F1/H4N7F1 | | X |
| H4N4E1F3 | | X |
| H4N6E1 | | X |
| H4N6E2 | | X |
| H4N6E2F1 | | X |
| H4N6E2F2 | | X |
| H4N7E2F1 | | X |
| H4N8Am1F3 | | X |
| H5N3E2F2 | | X |
| H5N4E1F3 | | X |
| H5N5E3 | | X |
| H5N5E3F1 | | X |
| H5N6Am2 | | X |
| H5N7Am1E2 | | X |
| H5N7Am1E2F1 | | X |
| H5N7E2 | | X |
| H5N7E3 | | X |
| H5N7E3F3 | | X |
| H6N3E2F1 | | X |
| H6N4Am1E3/H6N7E2 | | X |
| H6N4Am4 | | X |
| H6N4Am4F1 | | X |
| H6N4E3 | | X |
| H6N4E4 | | X |
| H7N4Am1E2F1/H7N7E1F1 | | X |
| H7N4E3F1 | | X |
| H7N4E4 | | X |
| H7N4E4F1 | | X |
| H7N5Am1E3/H7N8E2 | | X |
| H7N5Am2E1/H7N8Am1 | | X |
| H7N5Am4F1 | | X |
| H7N7E3F1 | | X |
| H7N8Am1E2 | | X |
| H7N8Am1E2F1 | | X |
| H7N8E3 | | X |
| H8N7Am2E1F1 | | X |
| H8N7Am3E1F2 | | X |
| H9N4Am3 | | X |
| H9N5 | | X |
| H10N2F1 | | |
| H10N3 | | |
| H10N3Am1 | | |
| H10N3Am1E1F1/H10N6F1 | | |
| H10N3Am1E1F2/H10N6F2 | | |
| H10N3Am1F1 | | |
| H10N3Am1F2 | | |
| H10N3Am2 | | |

| |
|---|
| H10N3Am2F1 |
| H10N3Am2F2 |
| H10N3E1 |
| H10N3E1F1 |
| H10N3E1F2 |
| H10N3E2 |
| H10N3E2F2 |
| H10N3F1 |
| H10N3F2 |
| H10N4 |
| H10N4Am1 |
| H10N4Am1E1/H10N7 |
| H10N4Am1E1F1/H10N7F1 |
| H10N4Am1E1F2/H10N7F2 |
| H10N4Am1E1F3/H10N7F3 |
| H10N4Am1E2/H10N7E1 |
| H10N4Am1E2F1/H10N7E1F1 |
| H10N4Am1E2F2/H10N7E1F2 |
| H10N4Am1E2F3/H10N7E1F3 |
| H10N4Am1E3/H10N7E2 |
| H10N4Am1E3F1/H10N7E2F1 |
| H10N4Am1E3F2/H10N7E2F2 |
| H10N4Am1E3F3/H10N7E2F3 |
| H10N4Am1F1 |
| H10N4Am1F2 |
| H10N4Am1F3 |
| H10N4Am2 |
| H10N4Am2E1/H10N7Am1 |
| H10N4Am2E1F1/H10N7Am1F1 |
| H10N4Am2E1F2/H10N7Am1F2 |
| H10N4Am2E1F3/H10N7Am1F3 |
| H10N4Am2E2/H10N7Am1E1 |
| H10N4Am2E2F1/H10N7Am1E1F1 |
| H10N4Am2E2F2/H10N7Am1E1F2 |
| H10N4Am2E2F3/H10N7Am1E1F3 |
| H10N4Am2F1 |
| H10N4Am2F2 |
| H10N4Am2F3 |
| H10N4Am3 |
| H10N4Am3E1/H10N7Am2 |
| H10N4Am3E1F1/H10N7Am2F1 |
| H10N4Am3E1F2/H10N7Am2F2 |
| H10N4Am3E1F3/H10N7Am2F3 |
| H10N4Am3F1 |
| H10N4Am3F2 |
| H10N4Am3F3 |
| H10N4Am4 |
| H10N4Am4F1 |
| H10N4Am4F2 |
| H10N4Am4F3 |

| |
|---|
| H10N4E1 |
| H10N4E1F1 |
| H10N4E1F2 |
| H10N4E1F3 |
| H10N4E2 |
| H10N4E2F1 |
| H10N4E2F2 |
| H10N4E2F3 |
| H10N4E3 |
| H10N4E3F1 |
| H10N4E3F2 |
| H10N4E3F3 |
| H10N4E4 |
| H10N4E4F1 |
| H10N4E4F2 |
| H10N4E4F3 |
| H10N4F1 |
| H10N4F2 |
| H10N4F3 |
| H10N5 |
| H10N5Am1E1F1/H10N8F1 |
| H10N5Am1E1F2/H10N8F2 |
| H10N5Am1E1F3/H10N8F3 |
| H10N5Am1E2/H10N8E1 |
| H10N5Am1E2F1/H10N8E1F1 |
| H10N5Am1E2F2/H10N8E1F2 |
| H10N5Am1E2F3/H10N8E1F3 |
| H10N5Am1E3/H10N8E2 |
| H10N5Am1E3F1/H10N8E2F1 |
| H10N5Am1E3F2/H10N8E2F2 |
| H10N5Am1E3F3 |
| H10N5Am1F1 |
| H10N5Am1F2 |
| H10N5Am1F3 |
| H10N5Am2E1/H10N8Am1 |
| H10N5Am2E1F1/H10N8Am1F1 |
| H10N5Am2E1F2/H10N8Am1F2 |
| H10N5Am2E1F3/H10N8Am1F3 |
| H10N5Am2E2/H10N8Am1E1 |
| H10N5Am2E2F1/H10N8Am1E1F1 |
| H10N5Am2E2F2/H10N8Am1E1F2 |
| H10N5Am2E2F3 |
| H10N5Am2F1 |
| H10N5Am2F2 |
| H10N5Am2F3 |
| H10N5Am3 |
| H10N5Am3E1/H10N8Am2 |
| H10N5Am3E1F1/H10N8Am2F1 |
| H10N5Am3E1F2/H10N8Am2F2 |
| H10N5Am3E1F3 |

| |
|---|
| H10N5Am3F1 |
| H10N5Am3F2 |
| H10N5Am3F3 |
| H10N5Am4 |
| H10N5Am4F1 |
| H10N5Am4F2 |
| H10N5Am4F3 |
| H10N5E1F1 |
| H10N5E1F2 |
| H10N5E1F3 |
| H10N5E2F1 |
| H10N5E2F2 |
| H10N5E2F3 |
| H10N5E3 |
| H10N5E3F1 |
| H10N5E3F2 |
| H10N5E3F3 |
| H10N5E4 |
| H10N5E4F1 |
| H10N5E4F2 |
| H10N5E4F3 |
| H10N5F2 |
| H10N5F3 |
| H10N6Am1 |
| H10N6Am1E1 |
| H10N6Am1E1F1 |
| H10N6Am1E1F2 |
| H10N6Am1E1F3 |
| H10N6Am1E2 |
| H10N6Am1E2F1 |
| H10N6Am1E2F2 |
| H10N6Am1E2F3 |
| H10N6Am1E3 |
| H10N6Am1E3F1 |
| H10N6Am1E3F2 |
| H10N6Am1F1 |
| H10N6Am1F2 |
| H10N6Am1F3 |
| H10N6Am2 |
| H10N6Am2E1 |
| H10N6Am2E1F1 |
| H10N6Am2E1F2 |
| H10N6Am2E1F3 |
| H10N6Am2E2 |
| H10N6Am2E2F1 |
| H10N6Am2E2F2 |
| H10N6Am2F1 |
| H10N6Am2F2 |
| H10N6Am2F3 |
| H10N6Am3 |

| |
|---|
| H10N6Am3E1 |
| H10N6Am3E1F1 |
| H10N6Am3E1F2 |
| H10N6Am3F1 |
| H10N6Am3F2 |
| H10N6Am3F3 |
| H10N6Am4 |
| H10N6Am4F1 |
| H10N6Am4F2 |
| H10N6E1 |
| H10N6E1F1 |
| H10N6E1F2 |
| H10N6E1F3 |
| H10N6E2 |
| H10N6E2F1 |
| H10N6E2F2 |
| H10N6E2F3 |
| H10N6E3 |
| H10N6E3F1 |
| H10N6E3F2 |
| H10N6E3F3 |
| H10N6E4 |
| H10N6E4F1 |
| H10N6E4F2 |
| H10N7Am1E2 |
| H10N7Am1E2F1 |
| H10N7Am1E2F2 |
| H10N7Am1E3 |
| H10N7Am1E3F1 |
| H10N7Am2E1 |
| H10N7Am2E1F1 |
| H10N7Am2E1F2 |
| H10N7Am2E2 |
| H10N7Am2E2F1 |
| H10N7Am3 |
| H10N7Am3E1 |
| H10N7Am3E1F1 |
| H10N7Am3F1 |
| H10N7Am3F2 |
| H10N7Am4 |
| H10N7Am4F1 |
| H10N7E3 |
| H10N7E3F1 |
| H10N7E3F2 |
| H10N7E4 |
| H10N7E4F1 |
| H10N8Am1E2 |
| H10N8Am1E2F1 |
| H10N8Am1E3 |
| H10N8Am2E1 |

| |
|---|
| H10N8Am2E1F1 |
| H10N8Am2E2 |
| H10N8Am3 |
| H10N8Am3E1 |
| H10N8Am3F1 |
| H10N8Am4 |
| H10N8E3 |
| H10N8E3F1 |
| H10N8E4 |
| H2N2 |
| H2N2F1 |
| H2N3 |
| H2N3F2 |
| H2N4 |
| H2N4F1 |
| H2N4F2 |
| H2N4F3 |
| H2N5 |
| H2N5F1 |
| H2N5F2 |
| H2N5F3 |
| H2N6 |
| H2N6F1 |
| H2N6F2 |
| H2N6F3 |
| H2N7 |
| H2N7F1 |
| H2N7F2 |
| H2N8 |
| H2N8F1 |
| H2N8F2 |
| H2N8F3 |
| H3N3E1F2 |
| H3N3F2 |
| H3N4Am1F2 |
| H3N4Am1F3 |
| H3N4E1F2 |
| H3N4E1F3 |
| H3N4F2 |
| H3N5Am1 |
| H3N5Am1F2 |
| H3N5Am1F3 |
| H3N5E1 |
| H3N5F2 |
| H3N5F3 |
| H3N6Am1 |
| H3N6Am1F2 |
| H3N6Am1F3 |
| H3N6E1 |
| H3N6E1F1 |

| |
|---|
| H3N6E1F2 |
| H3N6E1F3 |
| H3N6F2 |
| H3N6F3 |
| H3N7 |
| H3N7Am1 |
| H3N7Am1F2 |
| H3N7Am1F3 |
| H3N7E1 |
| H3N7E1F1 |
| H3N7E1F2 |
| H3N7E1F3 |
| H3N7F1 |
| H3N7F3 |
| H3N8 |
| H3N8Am1 |
| H3N8Am1F1 |
| H3N8Am1F2 |
| H3N8Am1F3 |
| H3N8E1 |
| H3N8E1F1 |
| H3N8E1F2 |
| H3N8E1F3 |
| H3N8F1 |
| H3N8F2 |
| H3N8F3 |
| H4N2F1 |
| H4N3Am1E1F2/H4N6F2 |
| H4N3Am2F1 |
| H4N3Am2F2 |
| H4N3E1F2 |
| H4N3E2F1 |
| H4N3E2F2 |
| H4N4Am1E1F3/H4N7F3 |
| H4N4Am1F3 |
| H4N4Am2F1 |
| H4N4Am2F2 |
| H4N4Am2F3 |
| H4N4E1F2 |
| H4N4E2F1 |
| H4N4E2F2 |
| H4N4E2F3 |
| H4N5Am1E1F1/H4N8F1 |
| H4N5Am1E1F3/H4N8F3 |
| H4N5Am1F3 |
| H4N5Am2F1 |
| H4N5Am2F3 |
| H4N5E1F3 |
| H4N5E2F1 |
| H4N5E2F3 |

| |
|---|
| H4N5F3 |
| H4N6Am1 |
| H4N6Am1E1F1 |
| H4N6Am1E1F2 |
| H4N6Am1E1F3 |
| H4N6Am1F1 |
| H4N6Am1F2 |
| H4N6Am1F3 |
| H4N6Am2 |
| H4N6Am2F1 |
| H4N6Am2F2 |
| H4N6Am2F3 |
| H4N6E1F2 |
| H4N6E1F3 |
| H4N6E2F3 |
| H4N6F3 |
| H4N7Am1 |
| H4N7Am1E1 |
| H4N7Am1E1F1 |
| H4N7Am1E1F2 |
| H4N7Am1F1 |
| H4N7Am1F2 |
| H4N7Am1F3 |
| H4N7Am2 |
| H4N7Am2F1 |
| H4N7Am2F2 |
| H4N7Am2F3 |
| H4N7E1F2 |
| H4N7E1F3 |
| H4N7E2 |
| H4N7E2F2 |
| H4N7E2F3 |
| H4N8Am1 |
| H4N8Am1E1 |
| H4N8Am1E1F1 |
| H4N8Am1E1F2 |
| H4N8Am1E1F3 |
| H4N8Am1F1 |
| H4N8Am1F2 |
| H4N8Am2 |
| H4N8Am2F1 |
| H4N8Am2F2 |
| H4N8Am2F3 |
| H4N8E1 |
| H4N8E1F1 |
| H4N8E1F2 |
| H4N8E1F3 |
| H4N8E2 |
| H4N8E2F1 |
| H4N8E2F2 |

| |
|---|
| H4N8E2F3 |
| H5N3Am1E1F1/H5N6F1 |
| H5N3Am1E1F2/H5N6F2 |
| H5N3Am1F2 |
| H5N3Am2F1 |
| H5N3Am2F2 |
| H5N3E1F2 |
| H5N3E2F1 |
| H5N3F2 |
| H5N4Am1E2/H5N7E1 |
| H5N4Am1E2F2/H5N7E1F2 |
| H5N4Am1E2F3/H5N7E1F3 |
| H5N4Am1F3 |
| H5N4Am2E1/H5N7Am1 |
| H5N4Am2E1F2/H5N7Am1F2 |
| H5N4Am2E1F3/H5N7Am1F3 |
| H5N4Am3 |
| H5N4Am3F1 |
| H5N4Am3F2 |
| H5N4Am3F3 |
| H5N4E3 |
| H5N4E3F1 |
| H5N4E3F2 |
| H5N4E3F3 |
| H5N5Am1E1F2/H5N8F2 |
| H5N5Am1E2/H5N8E1 |
| H5N5Am1E2F1/H5N8E1F1 |
| H5N5Am1E2F2/H5N8E1F2 |
| H5N5Am1E2F3/H5N8E1F3 |
| H5N5Am1F3 |
| H5N5Am2E1/H5N8Am1 |
| H5N5Am2E1F1/H5N8Am1F1 |
| H5N5Am2E1F2/H5N8Am1F2 |
| H5N5Am2E1F3/H5N8Am1F3 |
| H5N5Am2F2 |
| H5N5Am3 |
| H5N5Am3F1 |
| H5N5Am3F2 |
| H5N5Am3F3 |
| H5N5E1F3 |
| H5N5E2F2 |
| H5N5E3F2 |
| H5N5E3F3 |
| H5N5F3 |
| H5N6Am1E1 |
| H5N6Am1E1F2 |
| H5N6Am1E1F3 |
| H5N6Am1E2 |
| H5N6Am1E2F1 |
| H5N6Am1E2F2 |

| |
|---|
| H5N6Am1E2F3 |
| H5N6Am1F1 |
| H5N6Am1F2 |
| H5N6Am1F3 |
| H5N6Am2E1 |
| H5N6Am2E1F1 |
| H5N6Am2E1F2 |
| H5N6Am2E1F3 |
| H5N6Am2F2 |
| H5N6Am2F3 |
| H5N6Am3 |
| H5N6Am3F1 |
| H5N6Am3F2 |
| H5N6Am3F3 |
| H5N6E1F1 |
| H5N6E1F2 |
| H5N6E1F3 |
| H5N6E2 |
| H5N6E2F2 |
| H5N6E2F3 |
| H5N6E3 |
| H5N6E3F1 |
| H5N6E3F2 |
| H5N6E3F3 |
| H5N6F3 |
| H5N7Am1E1 |
| H5N7Am1E1F2 |
| H5N7Am1E1F3 |
| H5N7Am1E2F2 |
| H5N7Am1E2F3 |
| H5N7Am2 |
| H5N7Am2E1 |
| H5N7Am2E1F1 |
| H5N7Am2E1F2 |
| H5N7Am2E1F3 |
| H5N7Am2F2 |
| H5N7Am2F3 |
| H5N7Am3 |
| H5N7Am3F1 |
| H5N7Am3F2 |
| H5N7Am3F3 |
| H5N7E2F2 |
| H5N7E2F3 |
| H5N7E3F1 |
| H5N7E3F2 |
| H5N8Am1E1F2 |
| H5N8Am1E1F3 |
| H5N8Am1E2 |
| H5N8Am1E2F1 |
| H5N8Am1E2F2 |

| |
|---|
| H5N8Am1E2F3 |
| H5N8Am2 |
| H5N8Am2E1 |
| H5N8Am2E1F1 |
| H5N8Am2E1F2 |
| H5N8Am2E1F3 |
| H5N8Am2F1 |
| H5N8Am2F2 |
| H5N8Am2F3 |
| H5N8Am3 |
| H5N8Am3F1 |
| H5N8Am3F2 |
| H5N8Am3F3 |
| H5N8E2 |
| H5N8E2F1 |
| H5N8E2F2 |
| H5N8E2F3 |
| H5N8E3 |
| H5N8E3F1 |
| H5N8E3F2 |
| H5N8E3F3 |
| H6N2F1 |
| H6N3Am1E1F2/H6N6F2 |
| H6N3Am1F2 |
| H6N3Am2F1 |
| H6N3Am2F2 |
| H6N3E1F2 |
| H6N3E2F2 |
| H6N3F2 |
| H6N4Am1E1F2/H6N7F2 |
| H6N4Am1E1F3/H6N7F3 |
| H6N4Am1E2/H6N7E1 |
| H6N4Am1E2F1/H6N7E1F1 |
| H6N4Am1E2F2/H6N7E1F2 |
| H6N4Am1E2F3/H6N7E1F3 |
| H6N4Am1E3F1/H6N7E2F1 |
| H6N4Am1E3F2/H6N7E2F2 |
| H6N4Am1E3F3/H6N7E2F3 |
| H6N4Am1F2 |
| H6N4Am2E1/H6N7Am1 |
| H6N4Am2E1F1/H6N7Am1F1 |
| H6N4Am2E1F2/H6N7Am1F2 |
| H6N4Am2E1F3/H6N7Am1F3 |
| H6N4Am2E2/H6N7Am1E1 |
| H6N4Am2E2F1/H6N7Am1E1F1 |
| H6N4Am2E2F2/H6N7Am1E1F2 |
| H6N4Am2E2F3/H6N7Am1E1F3 |
| H6N4Am2F2 |
| H6N4Am2F3 |
| H6N4Am3 |

| |
|---|
| H6N4Am3E1/H6N7Am2 |
| H6N4Am3E1F1/H6N7Am2F1 |
| H6N4Am3E1F2/H6N7Am2F2 |
| H6N4Am3E1F3/H6N7Am2F3 |
| H6N4Am3F1 |
| H6N4Am3F2 |
| H6N4Am3F3 |
| H6N4Am4F2 |
| H6N4Am4F3 |
| H6N4E1F2 |
| H6N4E2F2 |
| H6N4E2F3 |
| H6N4E3F1 |
| H6N4E3F2 |
| H6N4E3F3 |
| H6N4E4F1 |
| H6N4E4F2 |
| H6N4E4F3 |
| H6N4F2 |
| H6N4F3 |
| H6N5Am1E1F3/H6N8F3 |
| H6N5Am1E2F3/H6N8E1F3 |
| H6N5Am1E3F1/H6N8E2F1 |
| H6N5Am1E3F2/H6N8E2F2 |
| H6N5Am1E3F3/H6N8E2F3 |
| H6N5Am1F3 |
| H6N5Am2E1F3/H6N8Am1F3 |
| H6N5Am2E2F1/H6N8Am1E1F1 |
| H6N5Am2E2F2/H6N8Am1E1F2 |
| H6N5Am2E2F3/H6N8Am1E1F3 |
| H6N5Am2F3 |
| H6N5Am3E1F1/H6N8Am2F1 |
| H6N5Am3E1F2/H6N8Am2F2 |
| H6N5Am3E1F3/H6N8Am2F3 |
| H6N5Am3F3 |
| H6N5Am4F1 |
| H6N5Am4F2 |
| H6N5Am4F3 |
| H6N5E1F3 |
| H6N5E2F3 |
| H6N5E3F3 |
| H6N5E4F1 |
| H6N5E4F2 |
| H6N5E4F3 |
| H6N5F3 |
| H6N6Am1E1F2 |
| H6N6Am1E1F3 |
| H6N6Am1E2F2 |
| H6N6Am1E2F3 |
| H6N6Am1E3 |

| |
|---|
| H6N6Am1E3F1 |
| H6N6Am1E3F2 |
| H6N6Am1E3F3 |
| H6N6Am2E1F3 |
| H6N6Am2E2 |
| H6N6Am2E2F1 |
| H6N6Am2E2F2 |
| H6N6Am2E2F3 |
| H6N6Am2F2 |
| H6N6Am2F3 |
| H6N6Am3E1 |
| H6N6Am3E1F1 |
| H6N6Am3E1F2 |
| H6N6Am3E1F3 |
| H6N6Am3F3 |
| H6N6Am4 |
| H6N6Am4F1 |
| H6N6Am4F2 |
| H6N6Am4F3 |
| H6N6E2F2 |
| H6N6E2F3 |
| H6N6E3F2 |
| H6N6E3F3 |
| H6N6E4 |
| H6N6E4F1 |
| H6N6E4F2 |
| H6N6E4F3 |
| H6N7Am1E2 |
| H6N7Am1E2F1 |
| H6N7Am1E2F2 |
| H6N7Am1E2F3 |
| H6N7Am1E3 |
| H6N7Am1E3F1 |
| H6N7Am1E3F2 |
| H6N7Am1E3F3 |
| H6N7Am2E1 |
| H6N7Am2E1F1 |
| H6N7Am2E1F2 |
| H6N7Am2E1F3 |
| H6N7Am2E2 |
| H6N7Am2E2F1 |
| H6N7Am2E2F2 |
| H6N7Am2E2F3 |
| H6N7Am3 |
| H6N7Am3E1 |
| H6N7Am3E1F1 |
| H6N7Am3E1F2 |
| H6N7Am3E1F3 |
| H6N7Am3F1 |
| H6N7Am3F2 |

| |
|---|
| H6N7Am3F3 |
| H6N7Am4 |
| H6N7Am4F1 |
| H6N7Am4F2 |
| H6N7Am4F3 |
| H6N7E3 |
| H6N7E3F1 |
| H6N7E3F2 |
| H6N7E3F3 |
| H6N7E4 |
| H6N7E4F1 |
| H6N7E4F2 |
| H6N7E4F3 |
| H6N8Am1E2 |
| H6N8Am1E2F1 |
| H6N8Am1E2F3 |
| H6N8Am1E3 |
| H6N8Am1E3F1 |
| H6N8Am1E3F2 |
| H6N8Am1E3F3 |
| H6N8Am2E1 |
| H6N8Am2E1F1 |
| H6N8Am2E1F3 |
| H6N8Am2E2 |
| H6N8Am2E2F1 |
| H6N8Am2E2F2 |
| H6N8Am2E2F3 |
| H6N8Am3 |
| H6N8Am3E1 |
| H6N8Am3E1F1 |
| H6N8Am3E1F2 |
| H6N8Am3E1F3 |
| H6N8Am3F1 |
| H6N8Am3F3 |
| H6N8Am4 |
| H6N8Am4F1 |
| H6N8Am4F2 |
| H6N8Am4F3 |
| H6N8E3 |
| H6N8E3F1 |
| H6N8E3F3 |
| H6N8E4 |
| H6N8E4F1 |
| H6N8E4F2 |
| H6N8E4F3 |
| H7N3 |
| H7N3Am1E1F1/H7N6F1 |
| H7N3Am1E1F2/H7N6F2 |
| H7N3Am2F1 |
| H7N3Am2F2 |

| |
|---|
| H7N3E2F1 |
| H7N3E2F2 |
| H7N4Am1E1/H7N7 |
| H7N4Am1E1F1/H7N7F1 |
| H7N4Am1E1F2/H7N7F2 |
| H7N4Am1E2/H7N7E1 |
| H7N4Am1E2F2/H7N7E1F2 |
| H7N4Am1E2F3/H7N7E1F3 |
| H7N4Am1E3/H7N7E2 |
| H7N4Am1E3F1/H7N7E2F1 |
| H7N4Am1E3F2/H7N7E2F2 |
| H7N4Am1E3F3/H7N7E2F3 |
| H7N4Am1F1 |
| H7N4Am1F2 |
| H7N4Am1F3 |
| H7N4Am2 |
| H7N4Am2E1/H7N7Am1 |
| H7N4Am2E1F1/H7N7Am1F1 |
| H7N4Am2E1F2/H7N7Am1F2 |
| H7N4Am2E1F3/H7N7Am1F3 |
| H7N4Am2E2/H7N7Am1E1 |
| H7N4Am2E2F1/H7N7Am1E1F1 |
| H7N4Am2E2F2/H7N7Am1E1F2 |
| H7N4Am2E2F3/H7N7Am1E1F3 |
| H7N4Am2F1 |
| H7N4Am2F2 |
| H7N4Am3 |
| H7N4Am3E1/H7N7Am2 |
| H7N4Am3E1F1/H7N7Am2F1 |
| H7N4Am3E1F2/H7N7Am2F2 |
| H7N4Am3E1F3/H7N7Am2F3 |
| H7N4Am3F1 |
| H7N4Am3F2 |
| H7N4Am3F3 |
| H7N4Am4 |
| H7N4Am4F1 |
| H7N4Am4F2 |
| H7N4Am4F3 |
| H7N4E1F1 |
| H7N4E1F2 |
| H7N4E1F3 |
| H7N4E2 |
| H7N4E2F1 |
| H7N4E2F2 |
| H7N4E3 |
| H7N4E3F2 |
| H7N4E3F3 |
| H7N4E4F2 |
| H7N4E4F3 |
| H7N4F1 |

| |
|---|
| H7N4F3 |
| H7N5 |
| H7N5Am1E1F2/H7N8F2 |
| H7N5Am1E2/H7N8E1 |
| H7N5Am1E2F2/H7N8E1F2 |
| H7N5Am1E2F3/H7N8E1F3 |
| H7N5Am1E3F1/H7N8E2F1 |
| H7N5Am1E3F2/H7N8E2F2 |
| H7N5Am1E3F3/H7N8E2F3 |
| H7N5Am1F2 |
| H7N5Am1F3 |
| H7N5Am2E1F2/H7N8Am1F2 |
| H7N5Am2E1F3/H7N8Am1F3 |
| H7N5Am2E2/H7N8Am1E1 |
| H7N5Am2E2F1/H7N8Am1E1F1 |
| H7N5Am2E2F2/H7N8Am1E1F2 |
| H7N5Am2E2F3/H7N8Am1E1F3 |
| H7N5Am2F2 |
| H7N5Am3 |
| H7N5Am3E1/H7N8Am2 |
| H7N5Am3E1F1/H7N8Am2F1 |
| H7N5Am3E1F2/H7N8Am2F2 |
| H7N5Am3E1F3/H7N8Am2F3 |
| H7N5Am3F2 |
| H7N5Am3F3 |
| H7N5Am4 |
| H7N5Am4F2 |
| H7N5Am4F3 |
| H7N5E1F2 |
| H7N5E1F3 |
| H7N5E2F2 |
| H7N5E3 |
| H7N5E3F2 |
| H7N5E3F3 |
| H7N5E4 |
| H7N5E4F1 |
| H7N5E4F2 |
| H7N5E4F3 |
| H7N5F2 |
| H7N5F3 |
| H7N6Am1E1F3 |
| H7N6Am1E2F3 |
| H7N6Am1E3F3 |
| H7N6Am2E1F3 |
| H7N6Am2E2F3 |
| H7N6Am2F3 |
| H7N6Am3F3 |
| H7N6E2F3 |
| H7N6E3F3 |
| H7N6E4F3 |

| |
|---|
| H7N6F3 |
| H7N7Am1E2F1 |
| H7N7Am1E2F2 |
| H7N7Am1E2F3 |
| H7N7Am1E3F1 |
| H7N7Am1E3F2 |
| H7N7Am1E3F3 |
| H7N7Am2E1F1 |
| H7N7Am2E1F2 |
| H7N7Am2E1F3 |
| H7N7Am2E2F1 |
| H7N7Am2E2F2 |
| H7N7Am2E2F3 |
| H7N7Am3E1F1 |
| H7N7Am3E1F2 |
| H7N7Am3E1F3 |
| H7N7Am3F1 |
| H7N7Am3F2 |
| H7N7Am3F3 |
| H7N7Am4F1 |
| H7N7Am4F2 |
| H7N7Am4F3 |
| H7N7E3F2 |
| H7N7E3F3 |
| H7N7E4F1 |
| H7N7E4F2 |
| H7N7E4F3 |
| H7N8Am1E2F2 |
| H7N8Am1E2F3 |
| H7N8Am1E3 |
| H7N8Am1E3F1 |
| H7N8Am1E3F2 |
| H7N8Am1E3F3 |
| H7N8Am2E1 |
| H7N8Am2E1F1 |
| H7N8Am2E1F2 |
| H7N8Am2E1F3 |
| H7N8Am2E2 |
| H7N8Am2E2F1 |
| H7N8Am2E2F2 |
| H7N8Am2E2F3 |
| H7N8Am3 |
| H7N8Am3E1 |
| H7N8Am3E1F1 |
| H7N8Am3E1F2 |
| H7N8Am3E1F3 |
| H7N8Am3F1 |
| H7N8Am3F2 |
| H7N8Am3F3 |
| H7N8Am4 |

| |
|---|
| H7N8Am4F1 |
| H7N8Am4F2 |
| H7N8Am4F3 |
| H7N8E3F1 |
| H7N8E3F2 |
| H7N8E3F3 |
| H7N8E4 |
| H7N8E4F1 |
| H7N8E4F2 |
| H7N8E4F3 |
| H8N2F1 |
| H8N3Am1E1F1/H8N6F1 |
| H8N3Am1F1 |
| H8N3Am1F2 |
| H8N3Am2F1 |
| H8N3Am2F2 |
| H8N3E1F1 |
| H8N3E1F2 |
| H8N3E2F1 |
| H8N3E2F2 |
| H8N3F2 |
| H8N4 |
| H8N4Am1 |
| H8N4Am1E1F3/H8N7F3 |
| H8N4Am1E2F1/H8N7E1F1 |
| H8N4Am1E2F2/H8N7E1F2 |
| H8N4Am1E2F3/H8N7E1F3 |
| H8N4Am1E3F2/H8N7E2F2 |
| H8N4Am1E3F3/H8N7E2F3 |
| H8N4Am1F1 |
| H8N4Am1F3 |
| H8N4Am2E1F1/H8N7Am1F1 |
| H8N4Am2E1F2/H8N7Am1F2 |
| H8N4Am2E1F3/H8N7Am1F3 |
| H8N4Am2E2F2/H8N7Am1E1F2 |
| H8N4Am2E2F3/H8N7Am1E1F3 |
| H8N4Am2F1 |
| H8N4Am2F3 |
| H8N4Am3 |
| H8N4Am3E1F2/H8N7Am2F2 |
| H8N4Am3E1F3/H8N7Am2F3 |
| H8N4Am3F1 |
| H8N4Am3F2 |
| H8N4Am3F3 |
| H8N4Am4 |
| H8N4Am4F1 |
| H8N4Am4F2 |
| H8N4Am4F3 |
| H8N4E1 |
| H8N4E1F1 |

| |
|---|
| H8N4E1F3 |
| H8N4E2F1 |
| H8N4E2F3 |
| H8N4E3 |
| H8N4E3F1 |
| H8N4E3F2 |
| H8N4E3F3 |
| H8N4E4 |
| H8N4E4F1 |
| H8N4E4F2 |
| H8N4E4F3 |
| H8N4F1 |
| H8N4F2 |
| H8N4F3 |
| H8N5Am1E1/H8N8 |
| H8N5Am1E1F1/H8N8F1 |
| H8N5Am1E1F3/H8N8F3 |
| H8N5Am1E2/H8N8E1 |
| H8N5Am1E2F1/H8N8E1F1 |
| H8N5Am1E2F2/H8N8E1F2 |
| H8N5Am1E2F3/H8N8E1F3 |
| H8N5Am1E3/H8N8E2 |
| H8N5Am1E3F1/H8N8E2F1 |
| H8N5Am1E3F2/H8N8E2F2 |
| H8N5Am1E3F3/H8N8E2F3 |
| H8N5Am2 |
| H8N5Am2E1/H8N8Am1 |
| H8N5Am2E1F1/H8N8Am1F1 |
| H8N5Am2E1F2/H8N8Am1F2 |
| H8N5Am2E1F3/H8N8Am1F3 |
| H8N5Am2E2/H8N8Am1E1 |
| H8N5Am2E2F1/H8N8Am1E1F1 |
| H8N5Am2E2F2/H8N8Am1E1F2 |
| H8N5Am2E2F3/H8N8Am1E1F3 |
| H8N5Am2F1 |
| H8N5Am2F2 |
| H8N5Am2F3 |
| H8N5Am3 |
| H8N5Am3E1/H8N8Am2 |
| H8N5Am3E1F1/H8N8Am2F1 |
| H8N5Am3E1F2/H8N8Am2F2 |
| H8N5Am3E1F3/H8N8Am2F3 |
| H8N5Am3F1 |
| H8N5Am3F2 |
| H8N5Am3F3 |
| H8N5Am4 |
| H8N5Am4F1 |
| H8N5Am4F2 |
| H8N5Am4F3 |
| H8N5E2 |

| |
|---|
| H8N5E2F1 |
| H8N5E2F2 |
| H8N5E2F3 |
| H8N5E3 |
| H8N5E3F1 |
| H8N5E3F2 |
| H8N5E3F3 |
| H8N5E4 |
| H8N5E4F1 |
| H8N5E4F2 |
| H8N5E4F3 |
| H8N5F1 |
| H8N5F3 |
| H8N6Am1 |
| H8N6Am1E1 |
| H8N6Am1E1F3 |
| H8N6Am1E2 |
| H8N6Am1E2F2 |
| H8N6Am1E3 |
| H8N6Am1E3F1 |
| H8N6Am1E3F2 |
| H8N6Am1E3F3 |
| H8N6Am1F2 |
| H8N6Am1F3 |
| H8N6Am2 |
| H8N6Am2E1 |
| H8N6Am2E1F2 |
| H8N6Am2E2 |
| H8N6Am2E2F1 |
| H8N6Am2E2F2 |
| H8N6Am2E2F3 |
| H8N6Am2F3 |
| H8N6Am3 |
| H8N6Am3E1 |
| H8N6Am3E1F1 |
| H8N6Am3E1F2 |
| H8N6Am3E1F3 |
| H8N6Am3F2 |
| H8N6Am4 |
| H8N6Am4F1 |
| H8N6Am4F2 |
| H8N6Am4F3 |
| H8N6E1 |
| H8N6E1F2 |
| H8N6E1F3 |
| H8N6E2 |
| H8N6E2F3 |
| H8N6E3 |
| H8N6E3F2 |
| H8N6E4 |

| |
|---|
| H8N6E4F1 |
| H8N6E4F2 |
| H8N6E4F3 |
| H8N6F3 |
| H8N7Am1E2F1 |
| H8N7Am1E3F2 |
| H8N7Am1E3F3 |
| H8N7Am2E2F3 |
| H8N7Am3 |
| H8N7Am3E1F3 |
| H8N7Am3F1 |
| H8N7Am4F2 |
| H8N7Am4F3 |
| H8N7E3F1 |
| H8N7E4F1 |
| H8N7E4F2 |
| H8N7E4F3 |
| H8N8Am1E2 |
| H8N8Am1E2F1 |
| H8N8Am1E2F2 |
| H8N8Am1E2F3 |
| H8N8Am1E3 |
| H8N8Am1E3F1 |
| H8N8Am1E3F2 |
| H8N8Am2E1 |
| H8N8Am2E1F1 |
| H8N8Am2E1F2 |
| H8N8Am2E1F3 |
| H8N8Am2E2 |
| H8N8Am2E2F1 |
| H8N8Am2E2F2 |
| H8N8Am3 |
| H8N8Am3E1 |
| H8N8Am3E1F1 |
| H8N8Am3E1F2 |
| H8N8Am3F1 |
| H8N8Am3F2 |
| H8N8Am3F3 |
| H8N8Am4 |
| H8N8Am4F1 |
| H8N8Am4F2 |
| H8N8E3 |
| H8N8E3F1 |
| H8N8E3F2 |
| H8N8E3F3 |
| H8N8E4 |
| H8N8E4F1 |
| H8N8E4F2 |
| H9N2F1 |
| H9N3 |

| |
|---|
| H9N3Am1 |
| H9N3Am1E1/H9N6 |
| H9N3Am1E1F1/H9N6F1 |
| H9N3Am1E1F2/H9N6F2 |
| H9N3Am1F1 |
| H9N3Am1F2 |
| H9N3Am2 |
| H9N3Am2F1 |
| H9N3Am2F2 |
| H9N3E1 |
| H9N3E1F1 |
| H9N3E1F2 |
| H9N3E2 |
| H9N3E2F1 |
| H9N3E2F2 |
| H9N3F1 |
| H9N4 |
| H9N4Am1E1/H9N7 |
| H9N4Am1E1F1/H9N7F1 |
| H9N4Am1E1F2/H9N7F2 |
| H9N4Am1E1F3/H9N7F3 |
| H9N4Am1E2F1/H9N7E1F1 |
| H9N4Am1E2F2/H9N7E1F2 |
| H9N4Am1E2F3/H9N7E1F3 |
| H9N4Am1E3/H9N7E2 |
| H9N4Am1E3F2/H9N7E2F2 |
| H9N4Am1E3F3/H9N7E2F3 |
| H9N4Am1F1 |
| H9N4Am1F2 |
| H9N4Am1F3 |
| H9N4Am2 |
| H9N4Am2E1F1/H9N7Am1F1 |
| H9N4Am2E1F2/H9N7Am1F2 |
| H9N4Am2E1F3/H9N7Am1F3 |
| H9N4Am2E2/H9N7Am1E1 |
| H9N4Am2E2F2/H9N7Am1E1F2 |
| H9N4Am2E2F3/H9N7Am1E1F3 |
| H9N4Am2F1 |
| H9N4Am2F2 |
| H9N4Am2F3 |
| H9N4Am3E1/H9N7Am2 |
| H9N4Am3E1F2/H9N7Am2F2 |
| H9N4Am3E1F3/H9N7Am2F3 |
| H9N4Am3F1 |
| H9N4Am3F2 |
| H9N4Am3F3 |
| H9N4Am4 |
| H9N4Am4F1 |
| H9N4Am4F2 |
| H9N4Am4F3 |

| |
|---|
| H9N4E1F1 |
| H9N4E1F2 |
| H9N4E1F3 |
| H9N4E2 |
| H9N4E2F1 |
| H9N4E2F2 |
| H9N4E2F3 |
| H9N4E3 |
| H9N4E3F1 |
| H9N4E3F2 |
| H9N4E3F3 |
| H9N4E4 |
| H9N4E4F1 |
| H9N4E4F2 |
| H9N4E4F3 |
| H9N4F1 |
| H9N4F2 |
| H9N4F3 |
| H9N5Am1 |
| H9N5Am1E1/H9N8 |
| H9N5Am1E1F2/H9N8F2 |
| H9N5Am1E1F3/H9N8F3 |
| H9N5Am1E2/H9N8E1 |
| H9N5Am1E2F1/H9N8E1F1 |
| H9N5Am1E2F2/H9N8E1F2 |
| H9N5Am1E3/H9N8E2 |
| H9N5Am1E3F1/H9N8E2F1 |
| H9N5Am1E3F2/H9N8E2F2 |
| H9N5Am1E3F3/H9N8E2F3 |
| H9N5Am1F1 |
| H9N5Am1F2 |
| H9N5Am1F3 |
| H9N5Am2 |
| H9N5Am2E1/H9N8Am1 |
| H9N5Am2E1F1/H9N8Am1F1 |
| H9N5Am2E1F2/H9N8Am1F2 |
| H9N5Am2E2/H9N8Am1E1 |
| H9N5Am2E2F1/H9N8Am1E1F1 |
| H9N5Am2E2F2/H9N8Am1E1F2 |
| H9N5Am2E2F3/H9N8Am1E1F3 |
| H9N5Am2F1 |
| H9N5Am2F2 |
| H9N5Am2F3 |
| H9N5Am3 |
| H9N5Am3E1/H9N8Am2 |
| H9N5Am3E1F1/H9N8Am2F1 |
| H9N5Am3E1F2/H9N8Am2F2 |
| H9N5Am3E1F3/H9N8Am2F3 |
| H9N5Am3F1 |
| H9N5Am3F2 |

| |
|---|
| H9N5Am3F3 |
| H9N5Am4 |
| H9N5Am4F1 |
| H9N5Am4F2 |
| H9N5Am4F3 |
| H9N5E1 |
| H9N5E1F1 |
| H9N5E1F2 |
| H9N5E1F3 |
| H9N5E2 |
| H9N5E2F1 |
| H9N5E2F2 |
| H9N5E2F3 |
| H9N5E3 |
| H9N5E3F1 |
| H9N5E3F2 |
| H9N5E3F3 |
| H9N5E4 |
| H9N5E4F1 |
| H9N5E4F2 |
| H9N5E4F3 |
| H9N5F1 |
| H9N5F2 |
| H9N5F3 |
| H9N6Am1E1F2 |
| H9N6Am1E1F3 |
| H9N6Am1E2 |
| H9N6Am1E2F1 |
| H9N6Am1E2F2 |
| H9N6Am1E2F3 |
| H9N6Am1E3 |
| H9N6Am1E3F1 |
| H9N6Am1E3F2 |
| H9N6Am1E3F3 |
| H9N6Am1F1 |
| H9N6Am1F3 |
| H9N6Am2E1 |
| H9N6Am2E1F1 |
| H9N6Am2E1F2 |
| H9N6Am2E1F3 |
| H9N6Am2E2 |
| H9N6Am2E2F1 |
| H9N6Am2E2F2 |
| H9N6Am2E2F3 |
| H9N6Am2F1 |
| H9N6Am2F2 |
| H9N6Am2F3 |
| H9N6Am3 |
| H9N6Am3E1 |
| H9N6Am3E1F1 |

| |
|---|
| H9N6Am3E1F2 |
| H9N6Am3E1F3 |
| H9N6Am3F1 |
| H9N6Am3F2 |
| H9N6Am3F3 |
| H9N6Am4 |
| H9N6Am4F1 |
| H9N6Am4F2 |
| H9N6Am4F3 |
| H9N6E1F1 |
| H9N6E1F3 |
| H9N6E2F1 |
| H9N6E2F2 |
| H9N6E2F3 |
| H9N6E3 |
| H9N6E3F1 |
| H9N6E3F2 |
| H9N6E3F3 |
| H9N6E4 |
| H9N6E4F1 |
| H9N6E4F2 |
| H9N6E4F3 |
| H9N6F3 |
| H9N7Am1E2 |
| H9N7Am1E2F1 |
| H9N7Am1E2F2 |
| H9N7Am1E2F3 |
| H9N7Am1E3 |
| H9N7Am1E3F1 |
| H9N7Am1E3F2 |
| H9N7Am2E1 |
| H9N7Am2E1F1 |
| H9N7Am2E1F2 |
| H9N7Am2E1F3 |
| H9N7Am2E2 |
| H9N7Am2E2F1 |
| H9N7Am2E2F2 |
| H9N7Am3 |
| H9N7Am3E1 |
| H9N7Am3E1F1 |
| H9N7Am3E1F2 |
| H9N7Am3F1 |
| H9N7Am3F2 |
| H9N7Am3F3 |
| H9N7Am4 |
| H9N7Am4F1 |
| H9N7Am4F2 |
| H9N7E3 |
| H9N7E3F1 |
| H9N7E3F2 |

| |
|---|
| H9N7E3F3 |
| H9N7E4 |
| H9N7E4F1 |
| H9N7E4F2 |
| H9N8Am1E2 |
| H9N8Am1E2F1 |
| H9N8Am1E2F2 |
| H9N8Am1E3 |
| H9N8Am1E3F1 |
| H9N8Am2E1F2 |
| H9N8Am2E2F1 |
| H9N8Am3 |
| H9N8Am3E1 |
| H9N8Am3E1F1 |
| H9N8Am3F1 |
| H9N8Am3F2 |
| H9N8Am4 |
| H9N8Am4F1 |
| H9N8E3 |
| H9N8E3F1 |
| H9N8E3F2 |
| H9N8E4 |
| H9N8E4F1 |

| Supplementary Table S-2. Identified O-glycan composition by original publication (Art. Id), GlycReSoft software tool (GlycReSoft Ids) and GlycoGenius (GG Ids). | | | |
|---|---|---|---|
| Glycan | GG Ids | Art. Ids | GlycReSoft Ids |
| H1F1 | X | X | X |
| H1N1 | X | X | X |
| N2 | X | X | X |
| H1X2 | X | X | X |
| H1N1F1 | X | X | X |
| H1N2 | X | X | X |
| H1N1S1 | X | X | X |
| H2N2 | X | X | X |
| H2N1S1 | X | X | X |
| H2N2F1 | X | X | X |
| H1N1S2 | X | X | X |
| H2N2S1 | X | X | X |
| H3N3 | X | X | X |
| H2N2S2 | X | X | X |
| H3N3S1 | X | X | X |
| H3N3S2 | X | X | X |
| H1N4F1 | X | | X |
| H3N2F2 | X | | X |
| H3N3F1 | X | | X |
| H4N3S1F1 | X | | X |
| H2N4S1F2 | X | | X |
| H5N4F1 | X | | X |
| H2N4S2F1 | X | | X |
| H3N4S1F2 | X | | X |
| H5N4S1 | X | | X |
| N1 | | X | X |
| H1X1 | | X | X |
| H1S1 | | X | X |
| H2N1 | | X | X |
| H2N1F1 | | X | X |
| H2N2S1F1 | | X | X |
| H2N3S1 | | X | X |
| H4N4S2 | | X | X |
| H2F1 | | X | |
| N1S1 | | X | |
| N2S1 | | X | |
| H1 | | | X |
| N1F1 | | | X |
| N1S1F1 | | | X |
| H3N2F1 | | | X |
| H3N2S1F1 | | | X |
| H3N2F3 | | | X |
| N2F1 | | | |
| H1S1F1 | | | |
| N3 | | | |
| H1N1F2 | | | |

| |
|---|
| N2F2 |
| H1N2F1 |
| N3F1 |
| H1N3 |
| H1N1S1F1 |
| H2N1F2 |
| N2S1F1 |
| H1N2S1 |
| H1N2F2 |
| H3N2 |
| N3S1 |
| N3F2 |
| H1N3F1 |
| H2N3 |
| H1N1S1F2 |
| H2N1S1F1 |
| H2N1F3 |
| H1N4 |
| N2S2 |
| N2S1F2 |
| H1N2S1F1 |
| H1N2F3 |
| H2N2F2 |
| N3S1F1 |
| N3F3 |
| H1N3S1 |
| H1N3F2 |
| H2N3F1 |
| H1N1S2F1 |
| H2N1S2 |
| H2N1S1F2 |
| N2S2F1 |
| H2N4 |
| H1N2S2 |
| H1N2S1F2 |
| H2N2F3 |
| H3N2S1 |
| N3S2 |
| N3S1F2 |
| H1N3S1F1 |
| H1N3F3 |
| H2N3F2 |
| H1N1S2F2 |
| H2N1S2F1 |
| H2N1S1F3 |
| H4N3 |
| H1N4S1 |
| H1N4F2 |
| N2S2F2 |
| H2N4F1 |

| |
|---|
| H1N2S2F1 |
| H1N2S1F3 |
| H3N4 |
| H2N2S1F2 |
| N3S2F1 |
| N3S1F3 |
| H2N5 |
| H1N3S2 |
| H1N3S1F2 |
| H2N3S1F1 |
| H2N3F3 |
| H3N3F2 |
| H2N1S3 |
| H2N1S2F2 |
| H4N3F1 |
| H1N4S1F1 |
| H1N4F3 |
| H2N4S1 |
| H2N4F2 |
| H1N2S3 |
| H1N2S2F2 |
| H3N4F1 |
| H2N2S2F1 |
| H2N2S1F3 |
| H4N4 |
| H3N2S2 |
| H3N2S1F2 |
| N3S3 |
| N3S2F2 |
| H2N5F1 |
| H1N3S2F1 |
| H1N3S1F3 |
| H3N5 |
| H2N3S2 |
| H2N3S1F2 |
| H3N3S1F1 |
| H3N3F3 |
| H2N1S3F1 |
| H2N1S2F3 |
| H4N3S1 |
| H4N3F2 |
| H1N4S2 |
| H1N4S1F2 |
| H2N4S1F1 |
| H2N4F3 |
| H1N2S3F1 |
| H1N2S2F3 |
| H3N4S1 |
| H3N4F2 |
| H2N2S3 |

| |
|---|
| H2N2S2F2 |
| H4N4F1 |
| H3N2S2F1 |
| H3N2S1F3 |
| H5N4 |
| N3S3F1 |
| N3S2F3 |
| H2N5S1 |
| H2N5F2 |
| H1N3S3 |
| H1N3S2F2 |
| H3N5F1 |
| H2N3S2F1 |
| H2N3S1F3 |
| H4N5 |
| H3N3S1F2 |
| H2N1S3F2 |
| H4N3F3 |
| H1N4S2F1 |
| H1N4S1F3 |
| H3N6 |
| H2N4S2 |
| H1N2S3F2 |
| H3N4S1F1 |
| H3N4F3 |
| H2N2S3F1 |
| H2N2S2F3 |
| H4N4S1 |
| H4N4F2 |
| H3N2S3 |
| H3N2S2F2 |
| N3S3F2 |
| H2N5S1F1 |
| H2N5F3 |
| H1N3S3F1 |
| H1N3S2F3 |
| H3N5S1 |
| H3N5F2 |
| H2N3S3 |
| H2N3S2F2 |
| H4N5F1 |
| H3N3S2F1 |
| H3N3S1F3 |
| H5N5 |
| H2N1S3F3 |
| H4N3S2 |
| H4N3S1F2 |
| H1N4S3 |
| H1N4S2F2 |
| H3N6F1 |

| |
|---|
| H2N4S1F3 |
| H4N6 |
| H1N2S3F3 |
| H3N4S2 |
| H2N2S4 |
| H2N2S3F2 |
| H4N4S1F1 |
| H3N2S3F1 |
| H3N2S2F3 |
| N3S3F3 |
| H2N5S2 |
| H2N5S1F2 |
| H1N3S4 |
| H1N3S3F2 |
| H3N5S1F1 |
| H2N3S3F1 |
| H2N3S2F3 |
| H4N5S1 |
| H3N3S3 |
| H3N3S2F2 |
| H4N3S2F1 |
| H1N4S3F1 |
| H1N4S2F3 |
| H3N6S1 |
| H2N4S3 |
| H2N4S2F2 |
| H3N4S2F1 |
| H2N2S4F1 |
| H2N2S3F3 |
| H3N2S4 |
| H3N2S3F2 |
| H2N5S2F1 |
| H1N3S4F1 |
| H1N3S3F3 |
| H3N5S2 |
| H2N3S4 |
| H2N3S3F2 |
| H3N3S3F1 |
| H4N3S3 |
| H1N4S4 |
| H1N4S3F2 |
| H2N4S3F1 |
| H3N4S3 |
| H2N2S4F2 |
| H3N2S4F1 |
| H2N5S3 |
| H1N3S4F2 |
| H2N3S4F1 |
| H3N3S4 |
| H1N4S4F1 |

| H2N4S4 |
|---|

**Supplementary Table S-3. Corrections calculated for fast isotopic distribution calculation.**

| Example Data | | All Atoms (Slow) | | | | Only Carbon (Fast) | | | | Ratio | | | | Corrected | | | | Differences(%) | | | | Actual Impact to Score(%) | | | |
|---|---|---|---|---|---|---|---|---|---|---|---|---|---|---|---|---|---|---|---|---|---|---|---|---|---|
| Glycan | Neutral Mass + Tag | 2nd Iso | 3rd Iso | 4th Iso | 5th Iso | 2nd Iso | 3rd Iso | 4th Iso | 5th Iso | 2nd Iso | 3rd Iso | 4th Iso | 5th Iso | 2nd Iso | 3rd Iso | 4th Iso | 5th Iso | 2nd Iso | 3rd Iso | 4th Iso | 5th Iso | 2nd Iso | 3rd Iso | 4th Iso | 5th Iso |
| H3N2 | 910.3278 | 0.375 | 0.122 | 0.028 | 0.006 | 0.368 | 0.066 | 0.008 | 0.001 | 1.019 | 1.848 | 3.500 | 6.000 | 0.375 | 0.115 | 0.025 | 0.005 | 0% | 6% | 11% | 17% | 0.0% | 1.2% | 0.6% | 0.3% |
| H4N2 | 1072.3806 | 0.440 | 0.158 | 0.041 | 0.009 | 0.433 | 0.091 | 0.013 | 0.001 | 1.016 | 1.736 | 3.154 | 9.000 | 0.441 | 0.153 | 0.039 | 0.008 | 0% | 3% | 5% | 11% | -0.2% | 0.6% | 0.3% | 0.2% |
| H5N2 | 1234.4334 | 0.505 | 0.199 | 0.057 | 0.014 | 0.498 | 0.121 | 0.019 | 0.002 | 1.014 | 1.645 | 3.000 | 7.000 | 0.507 | 0.195 | 0.055 | 0.013 | 0% | 2% | 4% | 7% | -0.3% | 0.4% | 0.2% | 0.1% |
| H3N3F1 | 1259.4651 | 0.530 | 0.210 | 0.061 | 0.015 | 0.519 | 0.132 | 0.022 | 0.003 | 1.021 | 1.591 | 2.773 | 5.000 | 0.530 | 0.212 | 0.062 | 0.015 | 0% | -1% | -2% | 0% | 0.0% | -0.2% | -0.1% | 0.0% |
| H6N2 | 1396.4863 | 0.570 | 0.243 | 0.077 | 0.021 | 0.562 | 0.155 | 0.028 | 0.004 | 1.014 | 1.568 | 2.750 | 5.250 | 0.574 | 0.242 | 0.075 | 0.019 | -1% | 0% | 3% | 10% | -0.5% | 0.1% | 0.2% | 0.2% |
| H3N3S1F1 | 1550.5605 | 0.653 | 0.298 | 0.102 | 0.029 | 0.638 | 0.200 | 0.041 | 0.006 | 1.024 | 1.490 | 2.488 | 4.833 | 0.651 | 0.304 | 0.104 | 0.030 | 0% | -2% | -2% | -3% | 0.2% | -0.4% | -0.1% | -0.1% |
| H7N2 | 1558.5391 | 0.635 | 0.292 | 0.100 | 0.029 | 0.627 | 0.193 | 0.039 | 0.006 | 1.013 | 1.513 | 2.564 | 4.833 | 0.640 | 0.293 | 0.099 | 0.028 | -1% | 0% | 1% | 3% | -0.6% | -0.1% | 0.1% | 0.1% |
| H3N3S1F2 | 1696.6184 | 0.718 | 0.350 | 0.128 | 0.039 | 0.703 | 0.243 | 0.055 | 0.009 | 1.021 | 1.440 | 2.327 | 4.333 | 0.717 | 0.361 | 0.134 | 0.041 | 0% | -3% | -5% | -5% | 0.1% | -0.6% | -0.3% | -0.1% |
| H8N2 | 1720.5919 | 0.700 | 0.346 | 0.128 | 0.040 | 0.692 | 0.236 | 0.053 | 0.009 | 1.012 | 1.466 | 2.415 | 4.444 | 0.706 | 0.348 | 0.126 | 0.038 | -1% | -1% | 2% | 5% | -0.6% | -0.1% | 0.1% | 0.1% |
| H9N2 | 1882.6447 | 0.764 | 0.403 | 0.159 | 0.053 | 0.757 | 0.283 | 0.069 | 0.013 | 1.009 | 1.424 | 2.304 | 4.077 | 0.772 | 0.407 | 0.158 | 0.051 | -1% | -1% | 1% | 4% | -0.7% | -0.2% | 0.0% | 0.1% |
| H5N4S1 | 1931.6876 | 0.808 | 0.433 | 0.174 | 0.058 | 0.790 | 0.307 | 0.079 | 0.015 | 1.023 | 1.410 | 2.203 | 3.867 | 0.805 | 0.440 | 0.178 | 0.060 | 0% | -2% | -2% | -3% | 0.3% | -0.3% | -0.1% | -0.1% |
| H7N4 | 1964.6978 | 0.815 | 0.443 | 0.180 | 0.061 | 0.800 | 0.316 | 0.082 | 0.016 | 1.019 | 1.402 | 2.195 | 3.813 | 0.816 | 0.451 | 0.183 | 0.062 | 0% | -2% | -2% | -2% | -0.1% | -0.4% | -0.1% | 0.0% |
| H5N4S2 | 2222.783 | 0.930 | 0.555 | 0.248 | 0.092 | 0.909 | 0.408 | 0.121 | 0.026 | 1.023 | 1.360 | 2.050 | 3.538 | 0.927 | 0.563 | 0.253 | 0.094 | 0% | -1% | -2% | -2% | 0.2% | -0.3% | -0.1% | 0.0% |
| H6N6F1 | 2354.8616 | 0.995 | 0.623 | 0.292 | 0.112 | 0.973 | 0.469 | 0.149 | 0.035 | 1.023 | 1.328 | 1.960 | 3.200 | 0.993 | 0.637 | 0.302 | 0.119 | 0% | -2% | -3% | -6% | 0.1% | -0.5% | -0.2% | -0.1% |
| H8N4S1 | 2417.8461 | 1.002 | 0.639 | 0.305 | 0.120 | 0.984 | 0.479 | 0.154 | 0.037 | 1.018 | 1.334 | 1.981 | 3.243 | 1.004 | 0.646 | 0.308 | 0.122 | 0% | -1% | -1% | -2% | -0.1% | -0.2% | -0.1% | 0.0% |
| H8N5S1 | 2620.9254 | 1.093 | 0.743 | 0.377 | 0.157 | 1.071 | 0.567 | 0.198 | 0.052 | 1.021 | 1.310 | 1.904 | 3.019 | 1.092 | 0.749 | 0.381 | 0.160 | 0% | -1% | -1% | -2% | 0.1% | -0.2% | -0.1% | 0.0% |
| H6N5S3 | 2879.0106 | 1.208 | 0.887 | 0.485 | 0.216 | 1.179 | 0.689 | 0.266 | 0.076 | 1.025 | 1.287 | 1.823 | 2.842 | 1.202 | 0.887 | 0.486 | 0.219 | 0% | 0% | 0% | -1% | 0.4% | 0.0% | 0.0% | 0.0% |
| H7N7S2 | 3156.1268 | 1.331 | 1.057 | 0.621 | 0.297 | 1.298 | 0.835 | 0.355 | 0.112 | 1.025 | 1.266 | 1.749 | 2.652 | 1.324 | 1.049 | 0.619 | 0.300 | 1% | 1% | 0% | -1% | 0.4% | 0.2% | 0.0% | 0.0% |
| H9N7F2 | 3190.1574 | 1.345 | 1.080 | 0.642 | 0.310 | 1.320 | 0.863 | 0.374 | 0.120 | 1.019 | 1.251 | 1.717 | 2.583 | 1.346 | 1.082 | 0.647 | 0.318 | 0% | 0% | -1% | -3% | -0.1% | 0.0% | 0.0% | 0.0% |
| H10N7S2 | 3642.2852 | 1.525 | 1.364 | 0.899 | 0.480 | 1.493 | 1.106 | 0.542 | 0.198 | 1.021 | 1.233 | 1.659 | 2.424 | 1.522 | 1.337 | 0.876 | 0.469 | 0% | 2% | 3% | 2% | 0.1% | 0.4% | 0.1% | 0.0% |
| H8N8S2F2 | 3813.3748 | 1.616 | 1.512 | 1.038 | 0.575 | 1.579 | 1.238 | 0.643 | 0.249 | 1.023 | 1.221 | 1.614 | 2.309 | 1.611 | 1.479 | 1.013 | 0.566 | 0% | 2% | 2% | 2% | 0.2% | 0.4% | 0.1% | 0.0% |
| H10N8S4 | 4427.5554 | 1.861 | 1.973 | 1.526 | 0.949 | 1.817 | 1.641 | 0.982 | 0.438 | 1.024 | 1.202 | 1.554 | 2.167 | 1.853 | 1.884 | 1.431 | 0.882 | 0% | 5% | 6% | 7% | 0.3% | 0.9% | 0.4% | 0.1% |
| | | | | | | | | | *Equations:* | *fast*1.02* | *fast*(10.8*(mass**-0.267))* | *fast*(122.62*(mass**-0.528))* | *fast*(2192.6*(mass**-0.833))* | | | | | | | | | | | | |

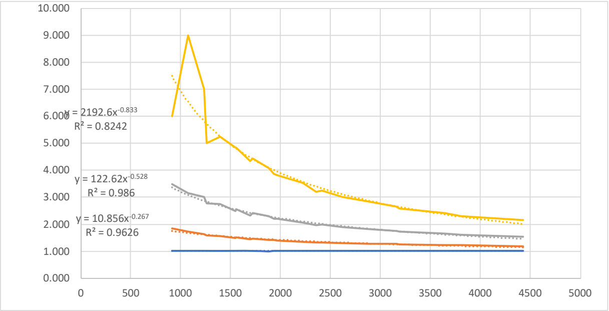

**Supplementary Table S-3. Model to estimate intensity of second isotopic envelope peak of a given molecule.**

| Fatty Acids | | *N* -Glycans | | Peptides | |
|---|---|---|---|---|---|
| ***mass*** | ***2nd peak*** | ***mass*** | ***2nd peak*** | ***mass*** | ***2nd peak*** |
| 74 | 0.034 | 1129.501 | 0.524 | 1390 | 0.672 |
| 130 | 0.079 | 1275.559 | 0.59 | 2749.333 | 1.31 |
| 200 | 0.134 | 1291.554 | 0.59 | 4168.997 | 2 |
| 340 | 0.246 | 1332.581 | 0.613 | 5558.663 | 2.66 |
| 761 | 0.58 | 1437.612 | 0.657 | 6948.329 | 3.39 |
| 2162 | 1.69 | 1453.607 | 0.657 | | |
| 2863 | 2.25 | 1478.639 | 0.68 | | |
| 3253 | 2.56 | 1494.634 | 0.68 | | |
| 3717 | 2.93 | 1535.66 | 0.702 | | |
| 4181 | 3.30 | 1599.665 | 0.724 | | |
| 4645 | 3.66 | 1615.66 | 0.724 | | |
| 5109 | 4.03 | 1624.697 | 0.746 | | |
| 5573 | 4.40 | 1640.691 | 0.746 | | |
| 6037 | 4.77 | 1656.686 | 0.746 | | |
| | | 1681.718 | 0.769 | | |
| | | 1697.713 | 0.769 | | |
| | | 1738.739 | 0.791 | | |
| | | 1761.718 | 0.791 | | |
| | | 1777.713 | 0.791 | | |
| | | 1785.729 | 0.802 | | |
| | | 1786.749 | 0.813 | | |
| | | 1802.744 | 0.813 | | |
| | | 1818.739 | 0.813 | | |
| | | 1827.776 | 0.836 | | |
| | | 1843.771 | 0.836 | | |
| | | 1859.766 | 0.836 | | |
| | | 1884.797 | 0.858 | | |
| | | 1900.792 | 0.858 | | |
| | | 1923.771 | 0.858 | | |
| | | 1931.787 | 0.869 | | |
| | | 1939.765 | 0.858 | | |
| | | 1941.819 | 0.88 | | |
| | | 1947.782 | 0.869 | | |
| | | 1948.802 | 0.88 | | |
| | | 1964.797 | 0.88 | | |
| | | 1980.792 | 0.88 | | |
| | | 1988.808 | 0.891 | | |
| | | 1989.829 | 0.902 | | |
| | | 2005.824 | 0.902 | | |
| | | 2021.819 | 0.902 | | |
| | | 2030.855 | 0.925 | | |
| | | 2046.85 | 0.925 | | |
| | | 2062.845 | 0.925 | | |
| | | 2077.845 | 0.936 | | |
| | | 2085.823 | 0.925 | | |
| | | 2087.877 | 0.947 | | |
| | | 2093.84 | 0.936 | | |
| | | 2101.818 | 0.925 | | |
| | | 2103.872 | 0.947 | | |
| | | 2109.835 | 0.936 | | |
| | | 2110.855 | 0.947 | | |
| | | 2126.85 | 0.947 | | |
| | | 2134.866 | 0.958 | | |
| | | 2142.845 | 0.947 | | |
| | | 2144.898 | 0.969 | | |

| Average | | | | |
|---|---|---|---|---|
| ***mass*** | ***2nd peak (lipids)*** | ***2nd peak (glycans)*** | ***2nd peak (proteins)*** | ***Average*** |
| 100 | 0.0555 | 0.507 | 0.0277 | 0.19673 |
| 200 | 0.1355 | 0.547 | 0.0777 | 0.2534 |
| 300 | 0.2155 | 0.587 | 0.1277 | 0.31007 |
| 400 | 0.2955 | 0.627 | 0.1777 | 0.36673 |
| 500 | 0.3755 | 0.667 | 0.2277 | 0.4234 |
| 600 | 0.4555 | 0.707 | 0.2777 | 0.48007 |
| 700 | 0.5355 | 0.747 | 0.3277 | 0.53673 |
| 800 | 0.6155 | 0.787 | 0.3777 | 0.5934 |
| 900 | 0.6955 | 0.827 | 0.4277 | 0.65007 |
| 1000 | 0.7755 | 0.867 | 0.4777 | 0.70673 |
| 1100 | 0.8555 | 0.907 | 0.5277 | 0.7634 |
| 1200 | 0.9355 | 0.947 | 0.5777 | 0.82007 |
| 1300 | 1.0155 | 0.987 | 0.6277 | 0.87673 |
| 1400 | 1.0955 | 1.027 | 0.6777 | 0.9334 |
| 1500 | 1.1755 | 1.067 | 0.7277 | 0.99007 |
| 1600 | 1.2555 | 1.107 | 0.7777 | 1.04673 |
| 1700 | 1.3355 | 1.147 | 0.8277 | 1.1034 |
| 1800 | 1.4155 | 1.187 | 0.8777 | 1.16007 |
| 1900 | 1.4955 | 1.227 | 0.9277 | 1.21673 |
| 2000 | 1.5755 | 1.267 | 0.9777 | 1.2734 |
| 2100 | 1.6555 | 1.307 | 1.0277 | 1.33007 |
| 2200 | 1.7355 | 1.347 | 1.0777 | 1.38673 |
| 2300 | 1.8155 | 1.387 | 1.1277 | 1.4434 |
| 2400 | 1.8955 | 1.427 | 1.1777 | 1.50007 |
| 2500 | 1.9755 | 1.467 | 1.2277 | 1.55673 |
| 2600 | 2.0555 | 1.507 | 1.2777 | 1.6134 |
| 2700 | 2.1355 | 1.547 | 1.3277 | 1.67007 |
| 2800 | 2.2155 | 1.587 | 1.3777 | 1.72673 |
| 2900 | 2.2955 | 1.627 | 1.4277 | 1.7834 |
| 3000 | 2.3755 | 1.667 | 1.4777 | 1.84007 |
| 3100 | 2.4555 | 1.707 | 1.5277 | 1.89673 |
| 3200 | 2.5355 | 1.747 | 1.5777 | 1.9534 |
| 3300 | 2.6155 | 1.787 | 1.6277 | 2.01007 |
| 3400 | 2.6955 | 1.827 | 1.6777 | 2.06673 |
| 3500 | 2.7755 | 1.867 | 1.7277 | 2.1234 |
| 3600 | 2.8555 | 1.907 | 1.7777 | 2.18007 |
| 3700 | 2.9355 | 1.947 | 1.8277 | 2.23673 |
| 3800 | 3.0155 | 1.987 | 1.8777 | 2.2934 |
| 3900 | 3.0955 | 2.027 | 1.9277 | 2.35007 |
| 4000 | 3.1755 | 2.067 | 1.9777 | 2.40673 |
| 4100 | 3.2555 | 2.107 | 2.0277 | 2.4634 |
| 4200 | 3.3355 | 2.147 | 2.0777 | 2.52007 |
| 4300 | 3.4155 | 2.187 | 2.1277 | 2.57673 |
| 4400 | 3.4955 | 2.227 | 2.1777 | 2.6334 |
| 4500 | 3.5755 | 2.267 | 2.2277 | 2.69007 |
| 4600 | 3.6555 | 2.307 | 2.2777 | 2.74673 |
| 4700 | 3.7355 | 2.347 | 2.3277 | 2.8034 |
| 4800 | 3.8155 | 2.387 | 2.3777 | 2.86007 |
| 4900 | 3.8955 | 2.427 | 2.4277 | 2.91673 |
| 5000 | 3.9755 | 2.467 | 2.4777 | 2.9734 |
| 5100 | 4.0555 | 2.507 | 2.5277 | 3.03007 |
| 5200 | 4.1355 | 2.547 | 2.5777 | 3.08673 |
| 5300 | 4.2155 | 2.587 | 2.6277 | 3.1434 |
| 5400 | 4.2955 | 2.627 | 2.6777 | 3.20007 |
| 5500 | 4.3755 | 2.667 | 2.7277 | 3.25673 |
| 5600 | 4.4555 | 2.707 | 2.7777 | 3.3134 |

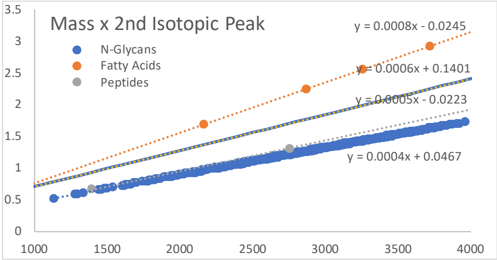

| | |
|---|---|
| 2150.861 | 0.958 |
| 2151.882 | 0.969 |
| 2167.877 | 0.969 |
| 2183.871 | 0.969 |
| 2191.888 | 0.98 |
| 2192.908 | 0.991 |
| 2208.903 | 0.991 |
| 2224.898 | 0.991 |
| 2233.935 | 1.014 |
| 2239.898 | 1.003 |
| 2247.876 | 0.991 |
| 2249.93 | 1.014 |
| 2255.893 | 1.003 |
| 2263.871 | 0.991 |
| 2265.925 | 1.014 |
| 2271.887 | 1.003 |
| 2272.908 | 1.014 |
| 2280.924 | 1.025 |
| 2288.903 | 1.014 |
| 2290.956 | 1.036 |
| 2296.919 | 1.025 |
| 2304.898 | 1.014 |
| 2306.951 | 1.036 |
| 2312.914 | 1.025 |
| 2313.934 | 1.036 |
| 2329.929 | 1.036 |
| 2337.946 | 1.047 |
| 2345.924 | 1.036 |
| 2347.978 | 1.058 |
| 2353.941 | 1.047 |
| 2354.961 | 1.058 |
| 2370.956 | 1.058 |
| 2386.951 | 1.058 |
| 2394.967 | 1.069 |
| 2395.988 | 1.081 |
| 2401.95 | 1.069 |
| 2409.929 | 1.058 |
| 2411.982 | 1.081 |
| 2417.945 | 1.069 |
| 2427.977 | 1.081 |
| 2433.94 | 1.069 |
| 2434.961 | 1.081 |
| 2437.014 | 1.103 |
| 2441.957 | 1.081 |
| 2442.977 | 1.092 |
| 2450.956 | 1.081 |
| 2453.009 | 1.103 |
| 2458.972 | 1.092 |
| 2466.951 | 1.081 |
| 2469.004 | 1.103 |
| 2474.967 | 1.092 |
| 2475.987 | 1.103 |
| 2484.004 | 1.114 |
| 2491.982 | 1.103 |
| 2494.036 | 1.125 |
| 2499.998 | 1.114 |
| 2507.977 | 1.103 |
| 2510.03 | 1.125 |
| 2515.993 | 1.114 |

| | | | | |
|---|---|---|---|---|
| 5700 | 4.5355 | 2.747 | 2.8277 | 3.37007 |
| 5800 | 4.6155 | 2.787 | 2.8777 | 3.42673 |
| 5900 | 4.6955 | 2.827 | 2.9277 | 3.4834 |
| 6000 | 4.7755 | 2.867 | 2.9777 | 3.54007 |
| 6100 | 4.8555 | 2.907 | 3.0277 | 3.59673 |
| 6200 | 4.9355 | 2.947 | 3.0777 | 3.6534 |
| 6300 | 5.0155 | 2.987 | 3.1277 | 3.71007 |
| 6400 | 5.0955 | 3.027 | 3.1777 | 3.76673 |
| 6500 | 5.1755 | 3.067 | 3.2277 | 3.8234 |
| 6600 | 5.2555 | 3.107 | 3.2777 | 3.88007 |
| 6700 | 5.3355 | 3.147 | 3.3277 | 3.93673 |
| 6800 | 5.4155 | 3.187 | 3.3777 | 3.9934 |
| 6900 | 5.4955 | 3.227 | 3.4277 | 4.05007 |
| 7000 | 5.5755 | 3.267 | 3.4777 | 4.10673 |
| 7100 | 5.6555 | 3.307 | 3.5277 | 4.1634 |
| 7200 | 5.7355 | 3.347 | 3.5777 | 4.22007 |
| 7300 | 5.8155 | 3.387 | 3.6277 | 4.27673 |
| 7400 | 5.8955 | 3.427 | 3.6777 | 4.3334 |
| 7500 | 5.9755 | 3.467 | 3.7277 | 4.39007 |
| 7600 | 6.0555 | 3.507 | 3.7777 | 4.44673 |
| 7700 | 6.1355 | 3.547 | 3.8277 | 4.5034 |
| 7800 | 6.2155 | 3.587 | 3.8777 | 4.56007 |
| 7900 | 6.2955 | 3.627 | 3.9277 | 4.61673 |
| 8000 | 6.3755 | 3.667 | 3.9777 | 4.6734 |
| 8100 | 6.4555 | 3.707 | 4.0277 | 4.73007 |
| 8200 | 6.5355 | 3.747 | 4.0777 | 4.78673 |
| 8300 | 6.6155 | 3.787 | 4.1277 | 4.8434 |
| 8400 | 6.6955 | 3.827 | 4.1777 | 4.90007 |
| 8500 | 6.7755 | 3.867 | 4.2277 | 4.95673 |
| 8600 | 6.8555 | 3.907 | 4.2777 | 5.0134 |
| 8700 | 6.9355 | 3.947 | 4.3277 | 5.07007 |
| 8800 | 7.0155 | 3.987 | 4.3777 | 5.12673 |
| 8900 | 7.0955 | 4.027 | 4.4277 | 5.1834 |
| 9000 | 7.1755 | 4.067 | 4.4777 | 5.24007 |
| 9100 | 7.2555 | 4.107 | 4.5277 | 5.29673 |
| 9200 | 7.3355 | 4.147 | 4.5777 | 5.3534 |
| 9300 | 7.4155 | 4.187 | 4.6277 | 5.41007 |
| 9400 | 7.4955 | 4.227 | 4.6777 | 5.46673 |
| 9500 | 7.5755 | 4.267 | 4.7277 | 5.5234 |
| 9600 | 7.6555 | 4.307 | 4.7777 | 5.58007 |
| 9700 | 7.7355 | 4.347 | 4.8277 | 5.63673 |
| 9800 | 7.8155 | 4.387 | 4.8777 | 5.6934 |
| 9900 | 7.8955 | 4.427 | 4.9277 | 5.75007 |
| 10000 | 7.9755 | 4.467 | 4.9777 | 5.80673 |

| | |
|---|---|
| 2517.014 | 1.125 |
| 2533.009 | 1.125 |
| 2541.025 | 1.136 |
| 2549.004 | 1.125 |
| 2557.02 | 1.136 |
| 2558.04 | 1.147 |
| 2564.003 | 1.136 |
| 2574.035 | 1.147 |
| 2579.998 | 1.136 |
| 2588.015 | 1.147 |
| 2590.03 | 1.147 |
| 2595.993 | 1.136 |
| 2597.014 | 1.147 |
| 2598.046 | 1.159 |
| 2599.067 | 1.17 |
| 2604.009 | 1.147 |
| 2605.03 | 1.159 |
| 2613.008 | 1.147 |
| 2615.062 | 1.17 |
| 2621.025 | 1.159 |
| 2631.057 | 1.17 |
| 2637.02 | 1.159 |
| 2638.04 | 1.17 |
| 2640.093 | 1.192 |
| 2645.036 | 1.17 |
| 2646.056 | 1.181 |
| 2654.035 | 1.17 |
| 2656.088 | 1.192 |
| 2662.051 | 1.181 |
| 2670.03 | 1.17 |
| 2672.083 | 1.192 |
| 2678.046 | 1.181 |
| 2679.067 | 1.192 |
| 2687.083 | 1.203 |
| 2695.062 | 1.192 |
| 2703.078 | 1.203 |
| 2711.056 | 1.192 |
| 2719.073 | 1.203 |
| 2720.093 | 1.214 |
| 2726.056 | 1.203 |
| 2734.072 | 1.214 |
| 2736.088 | 1.214 |
| 2742.051 | 1.203 |
| 2744.104 | 1.225 |
| 2750.067 | 1.214 |
| 2752.083 | 1.214 |
| 2758.046 | 1.203 |
| 2759.066 | 1.214 |
| 2760.099 | 1.225 |
| 2761.12 | 1.237 |
| 2766.062 | 1.214 |
| 2767.083 | 1.225 |
| 2777.115 | 1.237 |
| 2783.078 | 1.225 |
| 2791.094 | 1.237 |
| 2793.11 | 1.237 |
| 2799.072 | 1.225 |
| 2800.093 | 1.237 |
| 2801.126 | 1.248 |

| | |
|---|---|
| 2802.146 | 1.259 |
| 2807.089 | 1.237 |
| 2808.109 | 1.248 |
| 2816.088 | 1.237 |
| 2818.141 | 1.259 |
| 2824.104 | 1.248 |
| 2834.136 | 1.259 |
| 2840.099 | 1.248 |
| 2841.119 | 1.259 |
| 2848.115 | 1.259 |
| 2849.136 | 1.27 |
| 2857.114 | 1.259 |
| 2865.131 | 1.27 |
| 2873.109 | 1.259 |
| 2881.126 | 1.27 |
| 2882.146 | 1.281 |
| 2888.109 | 1.27 |
| 2890.162 | 1.292 |
| 2896.125 | 1.281 |
| 2898.141 | 1.281 |
| 2904.104 | 1.27 |
| 2906.157 | 1.292 |
| 2912.12 | 1.281 |
| 2914.136 | 1.281 |
| 2922.152 | 1.292 |
| 2923.173 | 1.303 |
| 2928.115 | 1.281 |
| 2929.135 | 1.292 |
| 2937.152 | 1.303 |
| 2939.167 | 1.303 |
| 2945.13 | 1.292 |
| 2947.184 | 1.315 |
| 2953.147 | 1.303 |
| 2955.162 | 1.303 |
| 2961.125 | 1.292 |
| 2962.146 | 1.303 |
| 2963.179 | 1.315 |
| 2964.199 | 1.326 |
| 2969.142 | 1.303 |
| 2970.162 | 1.315 |
| 2980.194 | 1.326 |
| 2986.157 | 1.315 |
| 2994.173 | 1.326 |
| 2996.189 | 1.326 |
| 3002.152 | 1.315 |
| 3003.172 | 1.326 |
| 3010.168 | 1.326 |
| 3011.189 | 1.337 |
| 3019.167 | 1.326 |
| 3027.183 | 1.337 |
| 3043.178 | 1.337 |
| 3044.199 | 1.348 |
| 3050.162 | 1.337 |
| 3051.195 | 1.348 |
| 3052.215 | 1.359 |
| 3058.178 | 1.348 |
| 3060.194 | 1.348 |
| 3068.21 | 1.359 |
| 3074.173 | 1.348 |

| | |
|---|---|
| 3076.189 | 1.348 |
| 3084.205 | 1.359 |
| 3085.225 | 1.37 |
| 3090.168 | 1.348 |
| 3091.188 | 1.359 |
| 3093.242 | 1.381 |
| 3098.184 | 1.359 |
| 3099.205 | 1.37 |
| 3101.22 | 1.37 |
| 3107.183 | 1.359 |
| 3109.237 | 1.381 |
| 3115.2 | 1.37 |
| 3117.215 | 1.37 |
| 3125.231 | 1.381 |
| 3126.252 | 1.393 |
| 3131.194 | 1.37 |
| 3132.215 | 1.381 |
| 3140.231 | 1.393 |
| 3142.247 | 1.393 |
| 3148.21 | 1.381 |
| 3156.226 | 1.393 |
| 3158.242 | 1.393 |
| 3164.205 | 1.381 |
| 3165.225 | 1.393 |
| 3172.221 | 1.393 |
| 3173.241 | 1.404 |
| 3189.236 | 1.404 |
| 3197.253 | 1.415 |
| 3205.231 | 1.404 |
| 3206.252 | 1.415 |
| 3213.248 | 1.415 |
| 3214.268 | 1.426 |
| 3220.231 | 1.415 |
| 3222.247 | 1.415 |
| 3230.263 | 1.426 |
| 3236.226 | 1.415 |
| 3244.242 | 1.426 |
| 3246.258 | 1.426 |
| 3247.278 | 1.437 |
| 3252.221 | 1.415 |
| 3253.241 | 1.426 |
| 3254.274 | 1.437 |
| 3255.294 | 1.448 |
| 3260.237 | 1.426 |
| 3261.257 | 1.437 |
| 3263.273 | 1.437 |
| 3271.289 | 1.448 |
| 3277.252 | 1.437 |
| 3279.268 | 1.437 |
| 3287.284 | 1.448 |
| 3288.305 | 1.459 |
| 3293.247 | 1.437 |
| 3294.268 | 1.448 |
| 3301.264 | 1.448 |
| 3302.284 | 1.459 |
| 3304.3 | 1.459 |
| 3310.263 | 1.448 |
| 3318.279 | 1.459 |
| 3320.295 | 1.459 |

| | |
|---|---|
| 3334.274 | 1.459 |
| 3335.294 | 1.471 |
| 3343.311 | 1.482 |
| 3351.289 | 1.471 |
| 3359.305 | 1.482 |
| 3367.284 | 1.471 |
| 3368.304 | 1.482 |
| 3375.3 | 1.482 |
| 3376.321 | 1.493 |
| 3382.284 | 1.482 |
| 3390.3 | 1.493 |
| 3392.316 | 1.493 |
| 3398.279 | 1.482 |
| 3400.332 | 1.504 |
| 3406.295 | 1.493 |
| 3408.311 | 1.493 |
| 3409.331 | 1.504 |
| 3416.327 | 1.504 |
| 3417.347 | 1.515 |
| 3422.29 | 1.493 |
| 3423.31 | 1.504 |
| 3425.326 | 1.504 |
| 3433.342 | 1.515 |
| 3439.305 | 1.504 |
| 3447.321 | 1.515 |
| 3449.337 | 1.515 |
| 3450.358 | 1.526 |
| 3455.3 | 1.504 |
| 3456.32 | 1.515 |
| 3463.316 | 1.515 |
| 3464.337 | 1.526 |
| 3466.352 | 1.526 |
| 3480.332 | 1.526 |
| 3482.347 | 1.526 |
| 3496.327 | 1.526 |
| 3497.347 | 1.537 |
| 3504.343 | 1.537 |
| 3505.363 | 1.548 |
| 3513.342 | 1.537 |
| 3521.358 | 1.548 |
| 3537.353 | 1.548 |
| 3538.374 | 1.56 |
| 3544.337 | 1.548 |
| 3546.39 | 1.571 |
| 3552.353 | 1.56 |
| 3554.369 | 1.56 |
| 3562.385 | 1.571 |
| 3568.348 | 1.56 |
| 3570.363 | 1.56 |
| 3578.38 | 1.571 |
| 3579.4 | 1.582 |
| 3584.343 | 1.56 |
| 3585.363 | 1.571 |
| 3593.379 | 1.582 |
| 3595.395 | 1.582 |
| 3601.358 | 1.571 |
| 3609.374 | 1.582 |
| 3611.39 | 1.582 |
| 3625.369 | 1.582 |

| | |
|---|---|
| 3626.39 | 1.593 |
| 3642.385 | 1.593 |
| 3650.401 | 1.604 |
| 3658.379 | 1.593 |
| 3666.396 | 1.604 |
| 3667.416 | 1.615 |
| 3683.411 | 1.615 |
| 3699.406 | 1.615 |
| 3707.422 | 1.626 |
| 3708.443 | 1.638 |
| 3714.406 | 1.626 |
| 3724.438 | 1.638 |
| 3730.401 | 1.626 |
| 3740.433 | 1.638 |
| 3746.396 | 1.626 |
| 3754.412 | 1.638 |
| 3755.432 | 1.649 |
| 3771.427 | 1.649 |
| 3787.422 | 1.649 |
| 3796.459 | 1.671 |
| 3812.454 | 1.671 |
| 3828.449 | 1.671 |
| 3853.48 | 1.693 |
| 3869.475 | 1.693 |
| 3900.47 | 1.704 |
| 3916.465 | 1.704 |
| 3957.491 | 1.727 |
